# The Virtual Plant Laboratory: a modern plant modeling framework in Julia

**DOI:** 10.1101/2024.09.27.615350

**Authors:** Alejandro Morales, David B. Kottelenberg, Ana Ernst, Rémi Vezy, Jochem B. Evers

## Abstract

The Virtual Plant Laboratory (VPL) is a novel software for building, simulating, and visualizing functional-structural plant (FSP) models. FSP models focus on the interactions between plant structure, internal physiological processes, and the biotic and abiotic environment. VPL is built in the Julia programming language and is designed to be a flexible and extensible platform for FSP modeling. Using Julia brings the advantage that only one programming language is required for the whole modeling cycle as Julia is as fast as compiled languages but also dynamic as interpreted languages. VPL provides a graph rewriting system for building dynamic models of plant growth and development, an interactive 3D visualization system and a Monte Carlo ray tracer for simulating radiation interception by plant canopies. In this paper, we introduce VPL, highlighting the main components, modeling paradigms, and design decisions behind it, as well as a future roadmap for further development. We also present a short case study of a model for intercropping of legumes and cereals that was built fully with VPL, as an example of what can be built with this software. VPL is fully open source and available in all common computing platforms for anyone to use. Full documentation and tutorials are available at https://virtualplantlab.com.

## Introduction

The Virtual Plant Laboratory (VPL) is a novel software for building, simulating, and visualizing functional-structural plant (FSP) models. FSP modeling focuses on the interactions between plant structure, internal physiological processes, and the biotic and abiotic environment (Buck-Sorlin, 2013, Godin and Sinoquet, 2005). FSP models span across multiple biological scales, from cells up to plant communities (Louarn and Song, 2020), and temporal scales, from calculations of instantaneous photosynthetic rates (Pearcy et al., 2005, Wang et al., 2017) to the growth and development throughout a plant’s life cycle (de Reffye et al., 2021). This makes FSP modeling an indispensable tool for (i) scaling up plant physiological processes, (ii) understanding the interplay between plant structure, physiology, and biotic and abiotic environment or (iii) addressing questions in ecology or plant production where the 3D structure and topology of plants play a significant role, in contrast to classic process-based models in which structure is not described explicitly (Buck-Sorlin, 2013).

In addition to covering multiple scales and addressing 3D plant structure explicitly, FSP models may also be interpreted as a special case of agent-based modeling (Zhang and DeAngelis, 2020). At each scale, FSP models can describe the different relevant structures (e.g., cells, organs, branches, plants) as separate entities, with an internal state and behavior. These entities interact with each other directly, through topological connections between organs or cells, or indirectly, through the environment. The dynamic interplay between direct and indirect interactions of these different entities results in emerging properties such as competition between plants for resources, which make FSP models a useful tool to explain complex spatial-temporal patterns in plant systems from simpler, local patterns.

These features that characterize FSP models also make them more challenging to implement than traditional mathematical or simulation models. For example, computing the interception of solar radiation by an explicit 3D structure requires using algorithms from computational geometry, many of which were originally developed for computer graphics, such as ray tracing (Bailey, 2018) or radiosity (Chelle and Andrieu, 1998). Similarly, keeping track of the topological relationships among organs of a plant over time requires using data structures, such as graphs (Godin and Caraglio, 1998, Hemmerling et al., 2008), whereas physiology and environmental physics are generally modeled by systems of algebraic and/or (partial) differential equations (Baldocchi et al., 2002, Farquhar et al., 1980). In contrast, a traditional process-based model of plant growth is often only comprised of a system of algebraic equations iterated over time (van Ittersum et al., 2003, Wang et al., 2002).

Given this need for multiple, non-trivial data structures and algorithms, specialized scientific software solutions (hereafter denoted as *modeling frameworks*) have been developed for FSP modeling, including GroIMP (Hemmerling et al., 2008), OpenAlea (Pradal et al., 2008) and L-Studio (Prusinkiewicz et al., 2000) among others (see below for details). These modeling frameworks allow modelers to focus on translating their scientific knowledge into code and execute it, without having to learn all the mathematical and computational details mentioned above. Modeling frameworks are fundamental to the recent growth in FSP modeling. Further progress in the field is, to some extent, limited by the improvement and maintenance of said frameworks. The Virtual Plant Laboratory (https://virtualplantlab.com) is a novel FSP modeling framework built in the Julia programming language aimed at allowing easier maintenance and further improvement.

In its current iteration, VPL focuses on FSP models defined at the levels of organ, plant and canopy (Buck-Sorlin, 2013). VPL is not intended to be used for models that describe the anatomy of organs (i.e., where individual cells or tissues are represented explicitly), such as existing 3D anatomical models of roots (Couvreur et al., 2018, Postma et al., 2017), leaves (Ho et al., 2016, Retta et al., 2023) or meristems (Smith et al., 2006). Similarly, VPL does not currently allow representing canopies with voxels or crown envelope volumes, which are often used when simulating trees (Chen et al., 2008, Duursma and Medlyn, 2012, Sinoquet et al., 2001).

In this paper, we first introduce the Virtual Plant Laboratory, highlighting the main components, modeling paradigms and design decisions behind it, as well as a future roadmap for further development. We also briefly describe the implementation of VPL in the Julia programming language. Finally, we present a short case study of a model for intercropping of legumes and cereals that was built with VPL, as an example of what can be built with this software. VPL is fully open source (all code is hosted in the VirtualPlantLab Github organization: https://github.com/VirtualPlantLab) and available in all common computing platforms for anyone to use (Linux, Windows, MacOS and FreeBSD). We refer potential users to the official website https://virtualplantlab.com for instructions on installation and use.

### The Virtual Plant Laboratory Modeling paradigms and features Plant topology (graphs)

VPL uses directed acyclic graphs, also known as tree graphs, to describe the topological description of a plant. Canonically, each node in a graph represents either an organ or a special topological feature that separates organs, such as the stem node that separates internodes, although this is not enforced. For instance, nodes can also be used to specify the origin and orientation of a plant or sub-plant scales, such as branches or tillers. A node in a graph may contain any data the user deems relevant, encapsulated within user-defined data types. Typically, these data will describe the geometry of an organ and its internal state like age, physiological signals, absorbed radiation, etc. It is also possible to store data at the graph level, which is intended to represent variables and parameters that are constant across organs or that are defined at the plant level, such as plant biomass. Constants and common parameters may also be defined as global variables outside of graphs.

In a dynamic FSP model, the graphs will evolve over time, either by changing the data stored in the nodes or by adding, subtracting and/or replacing nodes in the graph. To facilitate this dynamic evolution, VPL offers *rules* and *queries*. In both cases, a subset of the nodes in the graph will be selected based on a matching condition. This condition is defined by the type of node to be selected and a user-defined condition function that will be executed on each node of the right type, and which should return true or false. The condition function will have access to the data stored in the node, in the graph as well as all the other nodes in the graph through their topological relationships. The latter allows selecting nodes based on their context within the graph.

In the case of rules, the selected nodes will be replaced by a new node, a sub-graph (i.e., a set of connected nodes) or nothing, in which case the node and all the nodes downstream of it will be removed. The new node or sub-graph is created by a second user-defined function that has access to the same information as the condition function. This process of replacement of nodes is generally known as *graph rewriting* (Kurth et al., 2005) and multiple rules may be applied to a graph in an iterative fashion. These rules are conceptually defined to be executed in parallel to ensure that all rules apply to the same state of the graph. A consequence of this is that two rules cannot match the same node and all node matching must occur before any node is replaced. As a result, rules cannot be ordered, and must all be applied in every iteration.

In the case of queries, the selected nodes are returned in an array that the user can use to modify the internal state of those nodes. Queries do not rewrite the graph and they can be applied in any order and at any point within a simulation. Rules are used when changes to the topology of the plant are needed (e.g., emergence of new organs, pruning, senescence, etc.), whereas queries are used when only the internal state of existing nodes need to be adjusted (e.g., updating the dimensions of organs due to growth).

A graph in VPL is thus defined by three components: (i) an *axiom,* which is the initial set of nodes and their topological connections, (ii) a series of rules (if the model is dynamic) and, optionally, (iii) a user-defined object to store graph-level data. Queries are defined outside of graphs and are then applied to existing graphs to retrieve arrays of matched nodes.

VPL does not make any assumptions regarding time steps in the simulation. For example, it is possible to trigger the graph rewriting rules multiple times during a simulated day if the user wishes to do so. Generally, modifying the graphs over time thus consists of applications of rules, application of queries and modification of the internal states of the nodes matched by queries. The internal state of nodes may also be modified by the ray tracer (see below) in order to store absorbed radiation.

The graph rewriting paradigm described in the above can be considered to lie between the Lindenmayer systems of string rewriting or L-systems (Prusinkiewicz and Lindenmayer, 1990) and relational growth grammars (RGG, Hemmerling et al. 2008) However, a major departure from these paradigms is the fact that VPL does not use a grammar to define rules and queries but rather a procedural approach based on user-defined functions, data types and methods to traverse the graph and modify internal states, which relies on normal Julia syntax. This is by design, as we believe that such a procedural approach may be easier to learn and use by plant modelers without sacrificing generality. It also helps with debugging and testing as users have direct access to the code being executed (as opposed to grammars that generate the executable code for the user).

An exception to this procedural approach is the possibility in VPL to build sub-graphs for the axiom or the rules of a graph, using a simple algebra for graph construction. See Equation S2.1 for an example of a rule using the algebra for graph construction.

Finally, although VPL graphs do not encode hierarchical scales directly, nodes may represent subsets of the plant and hence graphs in VPL are compatible with the Multiscale Tree Graph formalism (MTG, Godin and Caraglio, 1998{, #7@@hidden}), the main difference being that the user would be responsible of keeping track of the different scales manually. Full compatibility with MTG, including the commonly used file format for data exchange, has not been incorporated yet but will be included in future releases of VPL.

### Plant structure (geometry)

VPL can convert a graph into a 3D representation for visualization and plant-environment simulation. The 3D representation consists of a series of triangular meshes, generated through a procedural geometry approach known as turtle graphics (Verhoeff, 2010). The *turtle* traverses the graph from the root node and execute user-defined methods to generate the geometry associated to each node in the graph. These methods are responsible for creating 3D meshes representing the structure of the plants, as well as color (for visualization, see below) and optical *materials* (for the ray tracer, see below). These methods can also affect the orientation and position of the turtle, i.e. its state, which is then kept for when it visits the next node in the graph. For simplicity, VPL offers multiple methods to modify the state of the turtle as well as generating the 3D meshes corresponding to common geometry primitives.

The turtle approach allows defining geometry locally, such as the angle of a leaf being specified relative to the branch it connects to rather than the horizontal plane, and the final geometry being generated will depend on the topological connections among nodes and the changes in the state of the turtle. This allows modelling complex plant architecture as an emergent property of simpler local geometry.

The generated 3D meshes alongside color and material objects are stored in a scene object which can then be used for ray tracing or visualization. In addition to generating these scenes from graphs, it is also possible to generate those 3D meshes directly and add them to an existing scene, alongside corresponding color and material objects. This is particularly useful when adding a soil layer to be able to simulate soil reflectance, or more complex environments such as the structure of a greenhouse compartment (Buck-Sorlin et al., 2011). VPL also allows exporting these 3D meshes into common file formats (e.g. PLY or OBJ) as well as importing them and we have successfully used exported 3D meshes of plants for 3D printing. Special methods are offered by VPL to scale and rotate imported meshes so that they can be used more easily in the context of a model.

One additional feature that has been included in VPL is the ability to generate different 3D meshes and/or color and material objects within the same simulation. Possible cases include: (i) different degree of geometric detail for visualization and ray tracing, (ii) removing some of the structure for specific visualizations, (iii) different ray tracers and therefore material objects or (iv) visualizing the same scene with different color schemes that map to properties stored in the nodes of the graphs. The latter allows 3D visualization to be used for the purpose of displaying organ properties like nitrogen content or absorbed radiation.

### Radiation interception (ray tracing)

In general, VPL only focuses on the algorithms and data structures that are specific to topology and structure in FSP modelling rather than on the functionality of said models, which the user can implement using a range of packages in Julia. However, the calculation of radiation interception by plants at the level of individual organs or below, is sufficiently complex that VPL comes with a solution for it. Several algorithms have been used in the past to model radiation interception by 3D plants, including ray tracing (Bailey, 2018, Cieslak et al., 2008, Wang et al., 2017), radiosity (Chelle and Andrieu, 1998) and rasterization (Vezy et al., 2020). In its current iteration, VPL offers a ray tracing algorithm.

The ray tracer within VPL allows for multiple wavelengths or wavebands to be simulated simultaneously, using the method proposed by Cieslak et al. (2008). VPL does not make any assumptions about which wavelengths are being simulated, nor the physical units in which radiation is expressed. Multiple radiation sources are available (point, line, area and directional) and several optical materials are offered (black, Lambertian, Phong and sensors that register radiation but do not affect it). New materials and radiation sources can be user-defined without modifying the source code of VPL, ensuring versatility.

The ray tracer may run in parallel across multiple cores, it is stochastic (different pseudo-random number generators may be used) and employs the Russian roulette mechanism proposed by Cieslak et al. (2008) to avoid errors in the energy balance when terminating rays. The intersection between rays and triangles is accelerated by creating a bounding volume hierarchy on the triangular meshes using surface area heuristic (MacDonald and Booth, 1990). This ensures that a minimum number of triangles need to be tested against each ray, reducing the overall computational cost.

Often, FSP models are approximating a canopy, whether growing in the field or under controlled conditions. In such cases, simulating hundreds of individual plant is often not feasible, so a smaller plot is typically modeled, although such downsizing can result in boundary effects when calculating radiation interception. To avoid this, the ray tracer of VPL implements a *grid cloner* based on the notion of graphics instancing (Marschner and Shirley, 2018), similar to the modelling framework GroIMP. This is an approach that emulates copying the scene a number of times along the different coordinate axes (horizontally or vertically). However, to avoid excessive memory consumption and computation from actually copying the meshes, graphics instancing will instead modify the origin of the ray being traced as if that ray was entering a neighboring clone of the scene. Note that the location of the virtual clones does not necessarily match the bounds of the scene, but rather it would be related to the spacing between plants. This means that the situation in which neighboring plants overlap is captured correctly by graphic instancing as opposed to, for example, periodic boundaries.

### Visualization (rendering)

VPL offers the possibility to render graphs (in 2D) and the 3D scenes generated from graphs and/or manually using an existing Julia package named Makie. For 3D scenes, it is also possible to render some of the elements used in the ray tracer, such as bounding boxes and radiation sources. VPL will return a data structure compatible with Makie that can be further modified by the user, so all the features in said package will be available for visualizations in VPL.

As of the time of writing, these backends include OpenGL, WebGL and Cairo. The latter can only generate static vector graphics so it is not useful for exploration of 3D scenes but it will work on headless systems that have no support for 3D graphics, and generally produce highest-quality plots for publications. The OpenGL and WebGL backends allow interacting with 3D scenes (rotation, panning and zooming) and 2D scenes (panning and zooming). WebGL is useful when running VPL in a web context such as Jupyter notebooks whereas OpenGL is the preferred option for local simulations. For OpenGL and WebGL backends, it is possible to export snapshots of the scene and create animations with scripted cameras.

Graph visualization will use a default node label (based on the type and internal node id) and color. However, the user can edit these properties for each type of node. In the case of 3D scenes, it is possible to visualize the wireframe pattern of the meshes (i.e., the edges of all the triangles) as well as the normal vectors for each triangle.

### Future roadmap

VPL is in constant development as new features are added to address the needs of the FSP modeling community. We do have the following roadmap which determines the current priorities for further development of VPL:

- Compatibility with Multiscale Tree Graphs (MTG). There is currently a Julia implementation of MTG (the MultiScaleTreeGraph.jl package (Vezy, 2023a)) which will serve as a starting point. Our aim is to be able to merge our current paradigm for graphs with the multiscale nature of MTG by adding new types of edges between nodes and adjusting the graph construction algebra. We expect this will make VPL more useful for simulations of larger, more complex plant structures (e.g., mature trees), especially when the simulation is initialized at an advanced stage of development. It may also be advantageous to have an explicit mechanism for storing data at different scales within the plant.
- Additional algorithms for radiation interception based on previous efforts. This is intended to reduce the computational cost of simulating the distribution of radiation within canopies in scenarios where the degree of realism that ray tracing ensures is not required. We note that the usual shortcuts that are used to reduce the computational burden of ray tracing (e.g., reduce the number of rays, coarse meshing of plant structure, coarse integration over the day, etc.) introduce numerical errors in the calculations of daily canopy photosynthesis (see Kent and Bailey (2021) for an example) in which cases faster, simpler approaches may be justified.
- Mapping graphs to systems of ordinary differential equations (ODE) for the purposes of simulating internal transport, such as for plant hydraulics, assimilate flow, hormonal signals, etc. Julia already offered state-of-the-art support for ODE systems, and there already examples of deriving ODE models from graphs, such as the package NetworkDynamics.jl (Lindner et al., 2021).
- Interface between the 3D structure of plants with environmental grids to simulate plant-environment interactions beyond radiation (e.g., temperature profiles within a canopy, soil water and heat transport, etc.). This would be achieved by *slicing* the scene along one or more axes and identifying the structural components in each of the resulting layers or voxels, above and/or belowground. This could then be used to aggregate fluxes and surface properties from the plants as inputs to models of environmental physics. This would lead to novel hybrid approaches, such as a soil-vegetation-transfer model where the vegetation is replaced by an FSP model.

### Implementation

VPL is implemented as a series of packages written entirely in Julia. In this section, we first justify why Julia is an appropriate choice for an FSP modeling framework. Then, we give an overview of how VPL is implemented in its current iteration, with an emphasis on the public API that users will make use when building models with VPL, as the internal details may change over time.

### Why Julia?

By using just-in-time compilation and a sophisticated system for code specialization, Julia achieves high performance while still being dynamic and highly interactive (Bezanson et al., 2017). That is, Julia breaks the “two language” paradigm that asserts one must choose between slow, dynamic, interactive languages (e.g., Python, R or Matlab) and fast, compiled, static languages (e.g., C, C++, Fortran or Java). In simple terms, Julia offers the convenience of Python but also the speed of C++.

By being implemented 100% in Julia and using common data structures, VPL is compatible with existing Julia packages to perform any modeling, simulation or visualization task beyond those specific to FSP modeling. For example, if the user needs to make use of a particular numerical method (e.g., ODE solver), import or export data or visualize the data generated by a simulation, they can choose any of the existing Julia packages to perform that task. Furthermore, Julia is highly interoperable with Python and R, so they can also use packages and code written in those languages from within their Julia scripts.

As VPL is written in Julia and supports multiple graphic backends, it is possible to execute VPL code in many different contexts such as (i) remote servers without graphics support, (ii) interactive computational notebooks such as Jupyter (Kluyver et al., 2016), Quarto (Allaire et al., 2022) or Pluto (van der Plas et al., 2023) (iii) IDEs that support Julia such as Visual Studio Code or (iv) simply by copying code into a terminal. Julia also supports Linux, Windows, MacOS and FreeBSD operating systems.

Julia and, by extension, VPL are also designed from the beginning for collaborative projects and code reuse. This implies favoring a decentralized, functional approach to programming, as opposed to the more traditional approach of building large software structures with complex class hierarchies. This philosophy has been followed in the development of VPL by (i) reusing existing Julia packages that implement specific functionalities and (ii) allowing users to modify the functionality of VPL as much as possible without touching the source code. Furthermore, we have split the implementation of VPL into multiple packages such that future collaborators can focus on specific aspects of FSP modelling. This approach to software design lowers the entry barrier for advanced users to become co-developers of VPL, favoring the longevity and sustainability of the software.

The internal structure of VPL and the lack of deep hierarchical classes, following Julia’s paradigm of multiple dispatch, makes it easier for users to define their own turtle commands, surface materials and even radiation sources, as they simply need to inherit from one abstract type and define a few methods per data type.

### Overview of VPL and the VPLverse

VPL is currently implemented as a Github organization comprised of:

- Five core packages that implement the basic functionality of the modelling framework (VPLcore)
- An interface package (VirtualPlantLab.jl) that exports the public API users are meant to use
- Four auxiliary packages that further aid in building and running FSP models (VPLverse)
- A collection of tutorials with examples of FSP models illustrating different features
- A website that collects all the documentation regarding VPL

The different repositories and packages are grouped into two sets (Figure 1). Firstly, the core packages plus interface that form VPL as described in the above. Secondly, auxiliary packages and repositories that complement VPL and are meant to be used with it, which we denote as the *VPLverse*.

**Figure 1.**
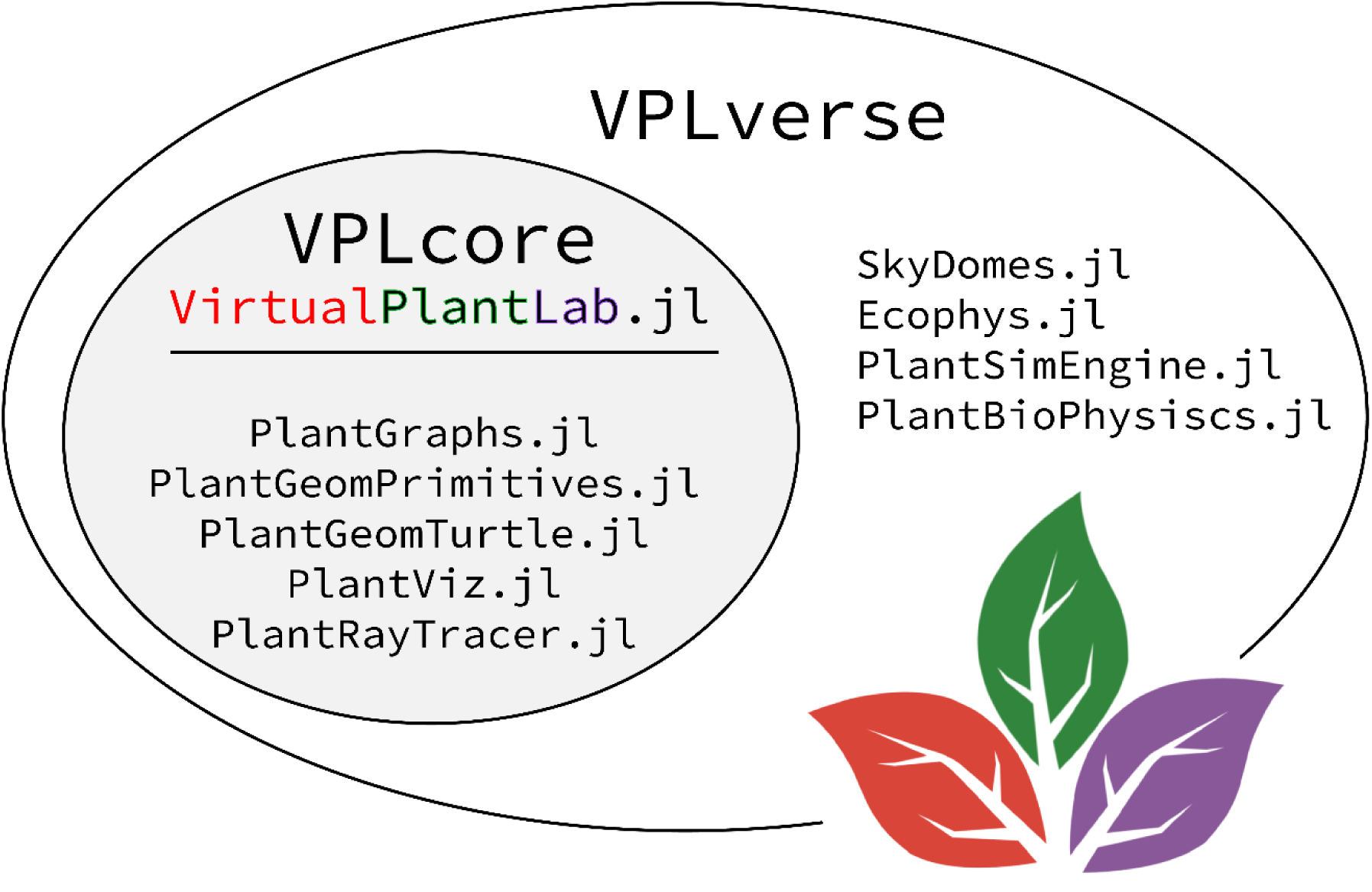
Organizational structure of VPL. The VPLcore includes the interface and 5 packages that integrate the basic functionality of VPL. Aside from the VPLcore, the VPLverse also includes other packages that are useful for the development of FSP models. All packages are suffixed with “.jl” as is customary in the Julia community.

The five core packages implement the basic data structures and algorithms required to build an FSP model:

- PlantGraphs.jl: A dynamic graph rewriting system where user-defined objects are stored in each node and these nodes can be queried and replaced by sub-graphs through dynamic production rules (see section on Plant Topology above).
- PlantGeomPrimitives.jl: A collection of 3D primitives implemented as triangular meshes stored in a scene for the purpose of visualization and ray tracing.
- PlantGeomTurtle.jl: An implementation of turtle algorithms that can generate 3D meshes from graphs representing the topology and structure of individual plants.
- PlantRayTracer.jl: A physics-based forward Monte Carlo ray tracer for the purpose of computing radiation interception by individual plants under field or controlled growth conditions. The ray tracer is multithreaded and uses bounding volume hierarchies.
- PlantViz.jl: 3D rendering of scenes based on the different backends of Makie.jl.

The VPLverse also comprises the following packages:

- SkyDomes.jl: generates radiation sources approximating the distribution of solar radiation in the sky (according to the CIE standards) as well as other variables related to solar geometry, daylength, etc.
- Ecophys.jl: Data structures and functions to simulate different ecophysiological processes, with an emphasis on short-term and long-term responses to aboveground environment (e.g., photosynthesis, transpiration, energy balance, phenology).
- PlantSimEngine.jl: Simulation engine for dynamic, process-based models (Vezy, 2023b). It handles model coupling, time stepping, and propagation of physical units or measurement errors through simulations. The package focuses on multiscale simulation and modeling of plants, soil, and atmosphere in interaction. It provides a user-friendly framework for defining processes and implementing associated models. With an emphasis on ease of use and high-performance computations, PlantSimEngine facilitates model coupling and modeling across different scales, from leaf-level simulations to entire scenes. Additionally, it supports various computation methods, including sequential, parallel, and distributed approaches.
- PlantBiophysics.jl: is a package to simulate biophysical processes for plants such as photosynthesis, conductance for heat, water vapor and CO₂, latent, sensible energy fluxes, net radiation and temperature. It leverages PlantSimEngine to declare the processes and implement the models, allowing for fast computations, *i.e.* just a few microseconds for the whole energy balance, photosynthesis and conductance coupling. This simulation speed makes it suitable for both model calibration and simulation tasks.

### Case study: Intercropping

To illustrate the possibilities of VPL, we implemented a model of an intercrop system composed of two species. This example illustrates the simulation of plants with different traits competing with each other for radiation, making use of most of the current components VPL has to offer.

Intercropping, the simultaneous cultivation of different crop species on the same land, has the potential to increase yield (Li et al., 2020b), water-use efficiency (Chen et al., 2018, Franco et al., 2018) and resource-use efficiency (Kermah et al., 2017, Tang et al., 2020), and reduce weed biomass (Gu et al., 2021) and pests (Li et al., 2021, Maitra, 2019) when compared to sole crops. A commonly employed intercropping system is cereal-legume row replacement intercropping, featuring alternating rows of cereals and legumes (Bedoussac et al., 2014, Kermah et al., 2017, Naudin et al., 2009) (Supplement 1 Figure S1.1).

Intercropping is an example of a heterogeneous canopy, featuring interactions among plants of the same and of different genotypes, leading to a variety of complex responses. This heterogeneity promotes canopy and root niche differentiation, potentially resulting in enhanced resource capture including radiation and soil nutrients (Brooker et al., 2015, Homulle et al., 2021, Li et al., 2020a). In response to these environmental cues, plants adapt their growth and developmental strategies to anticipate competition for resources. For instance, cereal plants adapt their elongation rate and tiller production in response to the red-to-far-red ratio (R:FR ratio), mitigating shading effects (Ballaré and Casal, 2000, Evers et al., 2006, Franklin, 2008). Choices in crop system design, such as intercropping versus sole cropping and plant density, therefore influence plant phenotypes.

FSP models are a valuable tool to dissect and analyze the intricate interactions between plants of different and the same crop species, and the impact of plant density (Evers et al., 2019, Gaudio et al., 2019). Therefore, for this case study, to illustrate the potential of the VPL platform, we developed an FSP model containing fictitious (but realistic) cereal and legume species with high and low-density settings. The full model description can be found in Supplement 2. The source code can be found inside the Gitlab servers within Wageningen University: https://git.wur.nl/david.kottelenberg/fspm_vpl_dk (in the branch named BASIC_CEREAL_LEGUME).

### Field setup

A cereal-legume row replacement intercrop, cereal sole crop, and legume sole crop were each simulated at two different plant densities (higher densities than usual practice were used to maximize plant-plant interactions). This resulted in six distinct simulation setups. Default densities were set at 480 plants m^−2^ for sole crop cereals and 128 plants m^−2^ for sole crop legumes, while high densities were set at 960 plants m^−2^ for sole crop cereals and 256 plants m^−2^ for sole crop legumes. Cereal-legume systems have alternating rows of cereals and legumes with the same density of plants in a row as the sole crop systems; i.e. the relative density of both crop species in the intercrop was 0.5. Row distances were uniformly set at 0.125 m and plot size remained constant at 2.0 x 0.5 m across all simulations. To mitigate border effects, plots were cloned five times in every cardinal direction, resulting in a field with 11 x 11 clones of the plot simulated. The grid cloner built into VPL was used for optimized performance.

### Simulations

Simulations were initiated on day 90 of the year and ran at a daily time step until the harvest date, set to 110 days after emergence (DAE) for both crops. Plants of both species emerged randomly on one of the first five simulation days. To account for model stochasticity, each simulation was run three times, resulting in a total of 18 simulations.

Air temperature was taken as the daily average temperature for each calendar day from the years 2001 to 2019 in De Bilt, The Netherlands (52° 06’ 36” N, 5° 10’ 50” E) (KNMI, 2024). Latitude for radiation intensity calculations was set at 52.0° N. Clear skies were simulated but we reduced incoming solar radiation by 40% in order to match the long-term average daily solar radiation in The Netherlands (Dalfsen, 1974). This resulted in an average photon flux of 640 µmol m^−2^ s^−1^ of direct radiation and 266 µmol m^−2^ s^−1^ of diffuse radiation per day over the whole growing season. In reality one would expect a higher proportion of diffuse radiation due to cloudiness, but our approximation is sufficient for the purpose of this case study.

### Output

To compare crop performance between the intercrop and sole crops, we calculated the biomass land equivalent ratio (LER) as the sum of partial LER (pLER) of each species where the pLER is the biomass of a species in the intercrop divided by the biomass of the same species in a monocrop setting (Li et al., 2023). See supplement 3 for equations regarding LER and pLER calculations. Typically, a pLER that is higher than the relative density of that species in the intercrop, is an indication for species overyielding in that intercrop. Thus, if LER >1, less land is needed to obtain the same total yields when intercropping compared to monocropping (even if only one of the species overyields).

The simulations outputs include aboveground crop biomass, plant height and R:FR ratios (see Supplement 2 for details on how these were calculated). Graphs were saved at every timestep, allowing revisitation of simulation days for additional data output. Subsequent data processing and visualization were done using R 4.3.2 (R Core Team, 2022) and the tidyverse package (Wickham et al., 2019).

### Simulation results

#### Biomass

Cereal plants produced more aboveground biomass in the intercrop compared to sole crop, particularly at the default density (Figure 2A). In contrast, legume plants produced more aboveground biomass within sole cropping systems than in the intercrop (Figure 2B). This indicates a competitive advantage of the cereal species over the legume species. Cereal plant biomass exhibited a multimodal distribution (Figure 2A), due to the presence of groups of high-performing plants, in contrast to the majority of lower-performing plants. Additionally, within the high-density intercrop, legume plants experienced substantial competitive pressure, leading to most plants having a biomass approaching zero (Figure 2B), indicating issues with plant survival.

**Figure 2.**
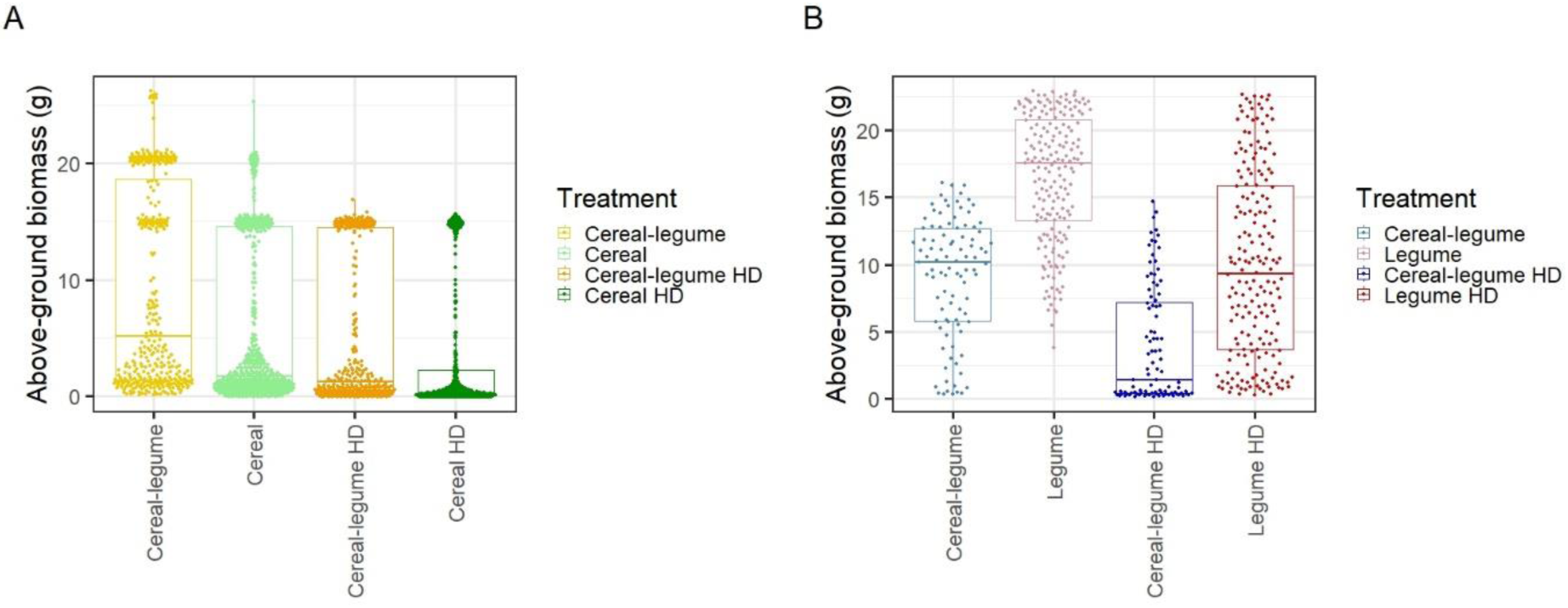
Box plots of plant biomass (in grams) across treatments by pooling the three replicates (as they do not vary much between them, see Supplement 3 Fig. S3.1), with individual plant biomass data represented by individual points. A: Cereal plant biomass. B: Legume plant biomass. HD: high density.

The intercrop at default density overproduced, with an LER of 1.12, while the high-density intercrop did not, with an LER of 0.99 (Figure 3). This indicates that the increase in cereal performance was higher than the decrease in legume performance at the standard density, but they compensated in the high-density intercrop.

**Figure 3.**
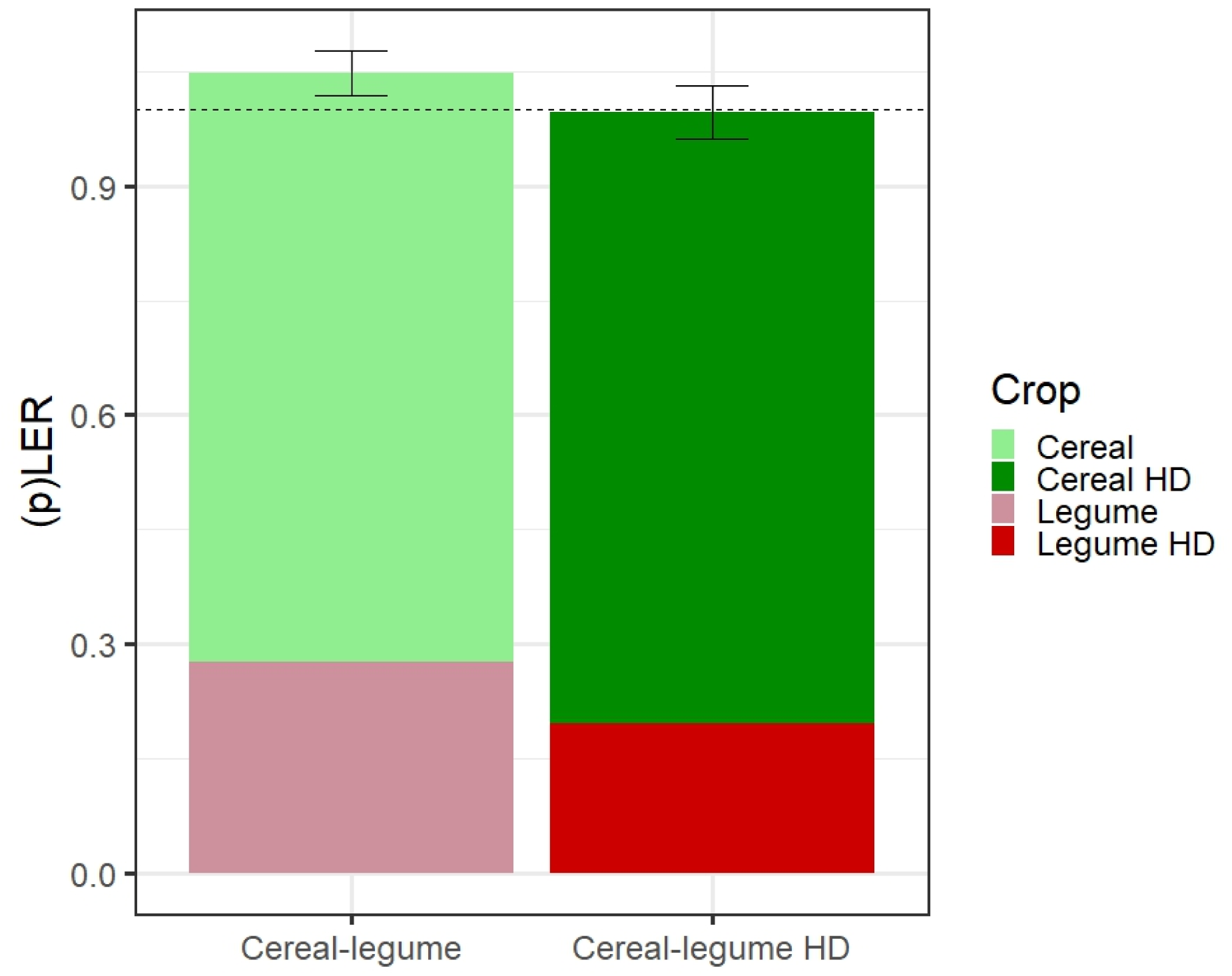
Bar plot of intercrop partial LERs (pLER) and the LER totals. Error bars indicate the standard deviation of the three replicates. HD: high density.

The difference in competitiveness is visually discernable at the individual plant level when coloring plant organs based on radiation capture (Figure 4). At 100 DAE, the canopy displayed a radiation capture gradient along the canopy height (Figure 4A). However, cereal plants appear brighter than legume plants, indicating that they captured more radiation, especially at the top of the plant (Figure 4B). To enhance visibility of the upper sections of legume plants, we only show two rows of the rendered field and filtered out cereal organs with a phytomer number exceeding six, counted from the bottom of the plant. This visual representation suggests that taller legume plants captured more radiation, evident in their brighter appearance at the plant’s top compared to shorter legume plants (Figure 4C). This nuanced visualization is possible due to the versatility provided by VPL for 3D rendering.

**Figure 4.**
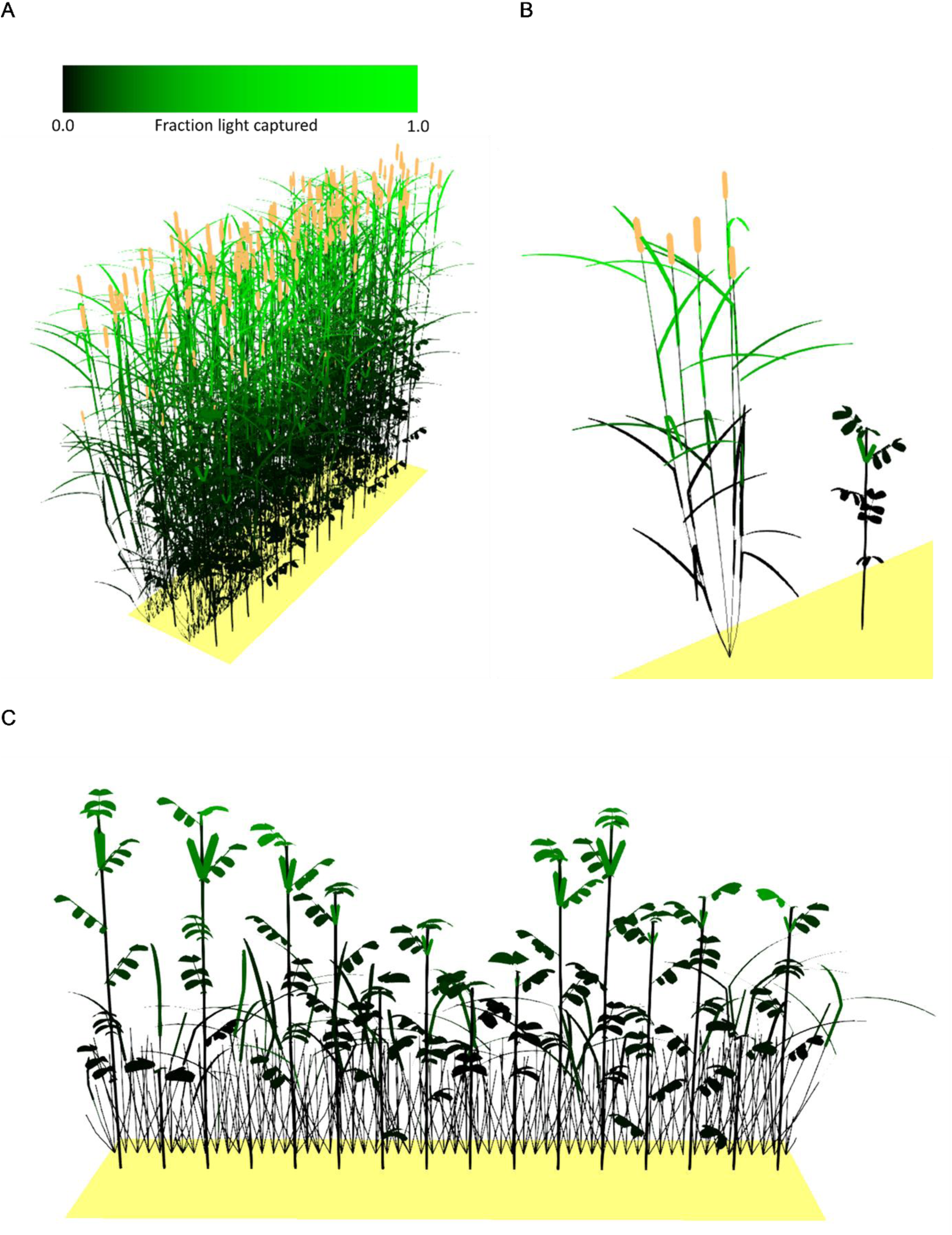
Render outputs with color based on fraction of radiation captured. A: Field render output of the normal density cereal-legume intercrop treatment at 100 DAE. B: Rendering of a single cereal and legume plant from the field in A. C: Partial view of the field in A, featuring only two rows, with cereal organs having a cumulative phytomer number exceeding six rendered transparent to enhance visibility of the upper sections of legume plants.

### Plant height

At the default density, plants were taller compared to high-density (Figure 5). Furthermore, plants were taller in the intercrop compared to sole crop. The influence of competition dynamics on plant height is mainly influenced by two aspects working in opposite directions and therefore displayed a nuanced response. In anticipation of competition, environmental signals can induce increase in height through an adaptive elongation response (Supplement 2, Equation S2.8). However, strong competition can reduce radiation capture by the plant, limiting resources available for elongation. Both aspects were contributing to the differences in height in cereal plants (Supplement Figure S3.2).

**Figure 5.**
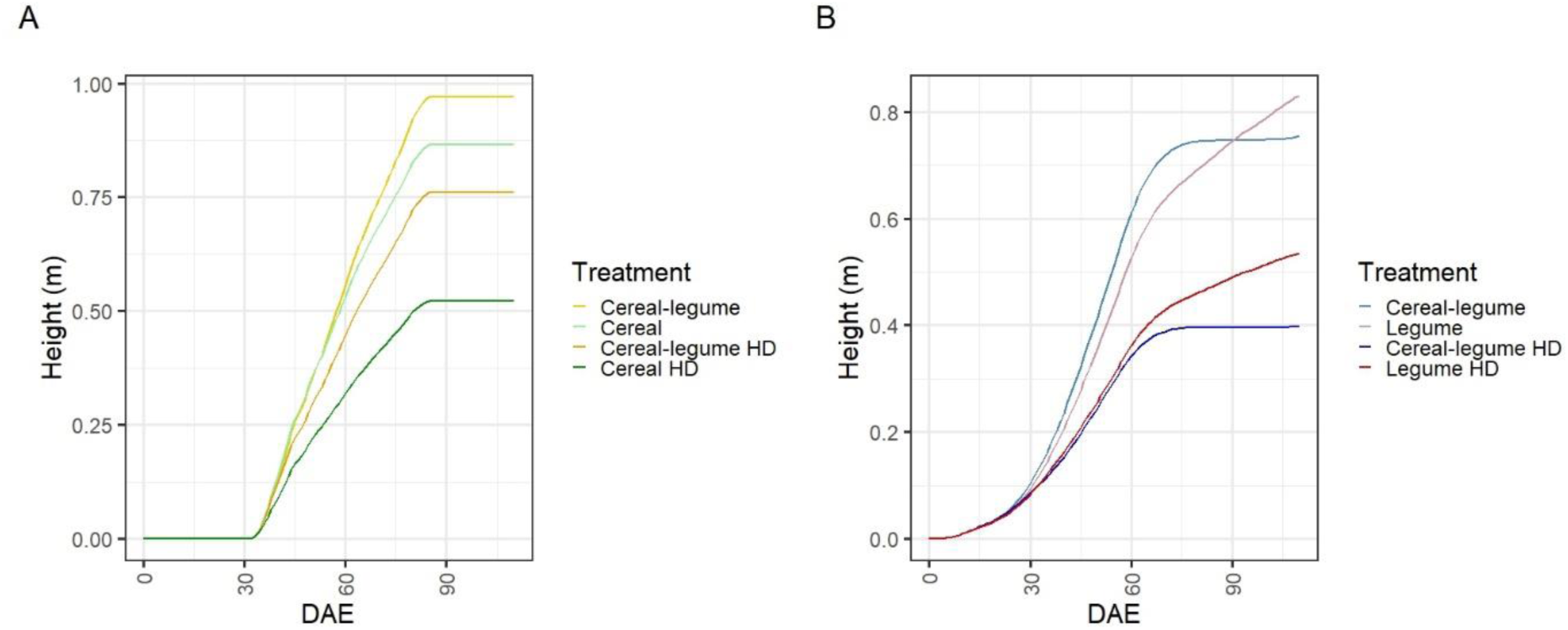
Plant height (in meters) over time in days after emergence (DAE). A: Cereal plant height across treatments. B: Legume plant height across treatments. HD: high density.

### Tillering

The number of tillers (i.e., branches of cereal plants) varied with level of competition, with increased competition resulting in fewer tillers (Figure 6A). The effect of the treatments on the number of tillers can be explained by the differences in detected R:FR ratio at the bottom of the canopy during the initial 30 DAE, as no more tillers were formed after (Figure 6B).

**Figure 6.**
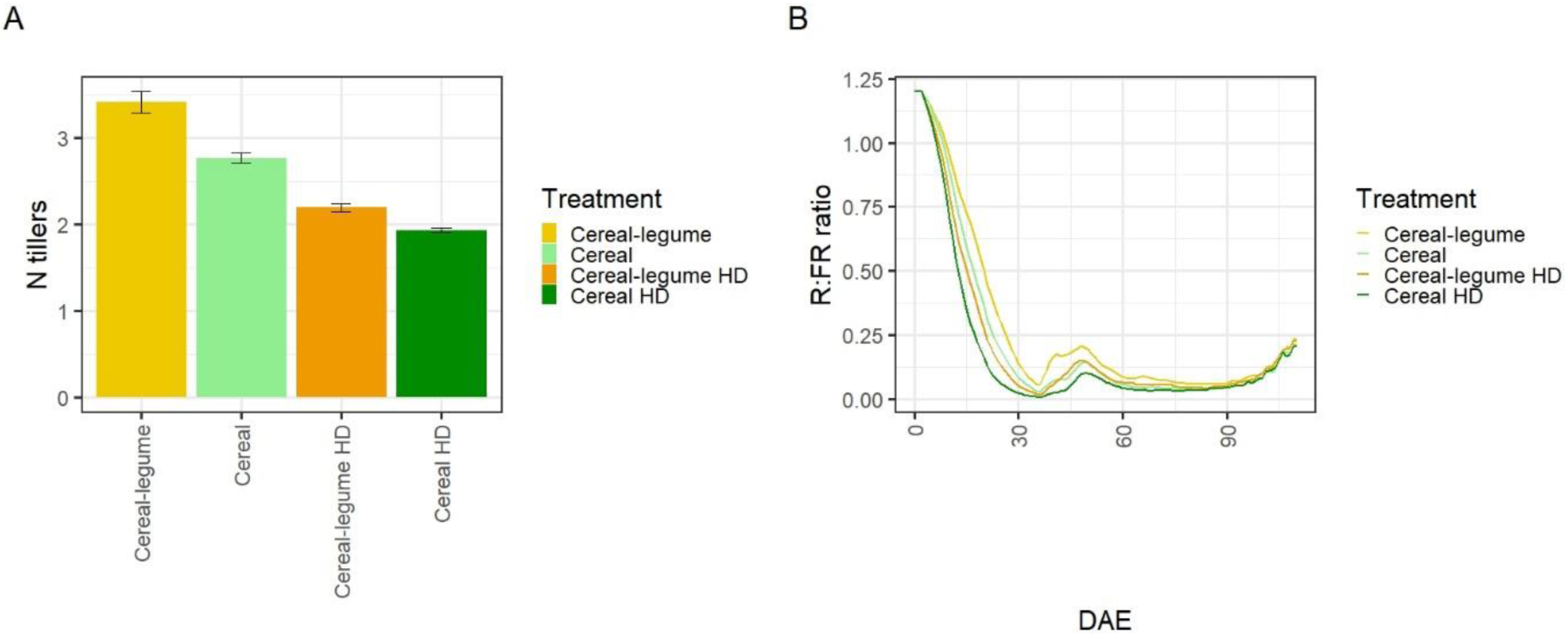
A: Number of cereal plant tillers created across treatments at 35 DAE. Error bars show the standard deviation of the three replicates. B: Red:far-red (R:FR) ratio detected at the bottom of cereal plants across treatments over time in days after emergence (DAE). HD: high density.

Although the R:FR ratio detected by a plant is similar to the proportion of radiation captured by a plant, a lower R:FR ratio can be observed earlier than a drop in radiation capture, as reduced-R:FR radiation reflected off neighboring plants is detected before shading takes place. Therefore, R:FR serves as an early signal of potential future shading due to the high reflection of far-red radiation by neighboring plants. This can be visualized in VPL by comparing R:FR ratio-based coloration of plant organs at 40 DAE between treatments with high and low competition, e.g. the cereal-legume treatment (Figure 7A) and high-density cereal sole crop treatment (Figure 7B). With heightened competition, the cereal plant shows darker-colored organs compared to the one experiencing lower competition. To emphasize this, restricting the visualization to organs with an R:FR ratio < 0.4 reveals the absence of leaves in the plant experiencing lower competition (Figure 7C), in contrast to the plant with higher competition (Figure 7D). This disparity underscores the impact of environmental signals on plant responses.

**Figure 7.**
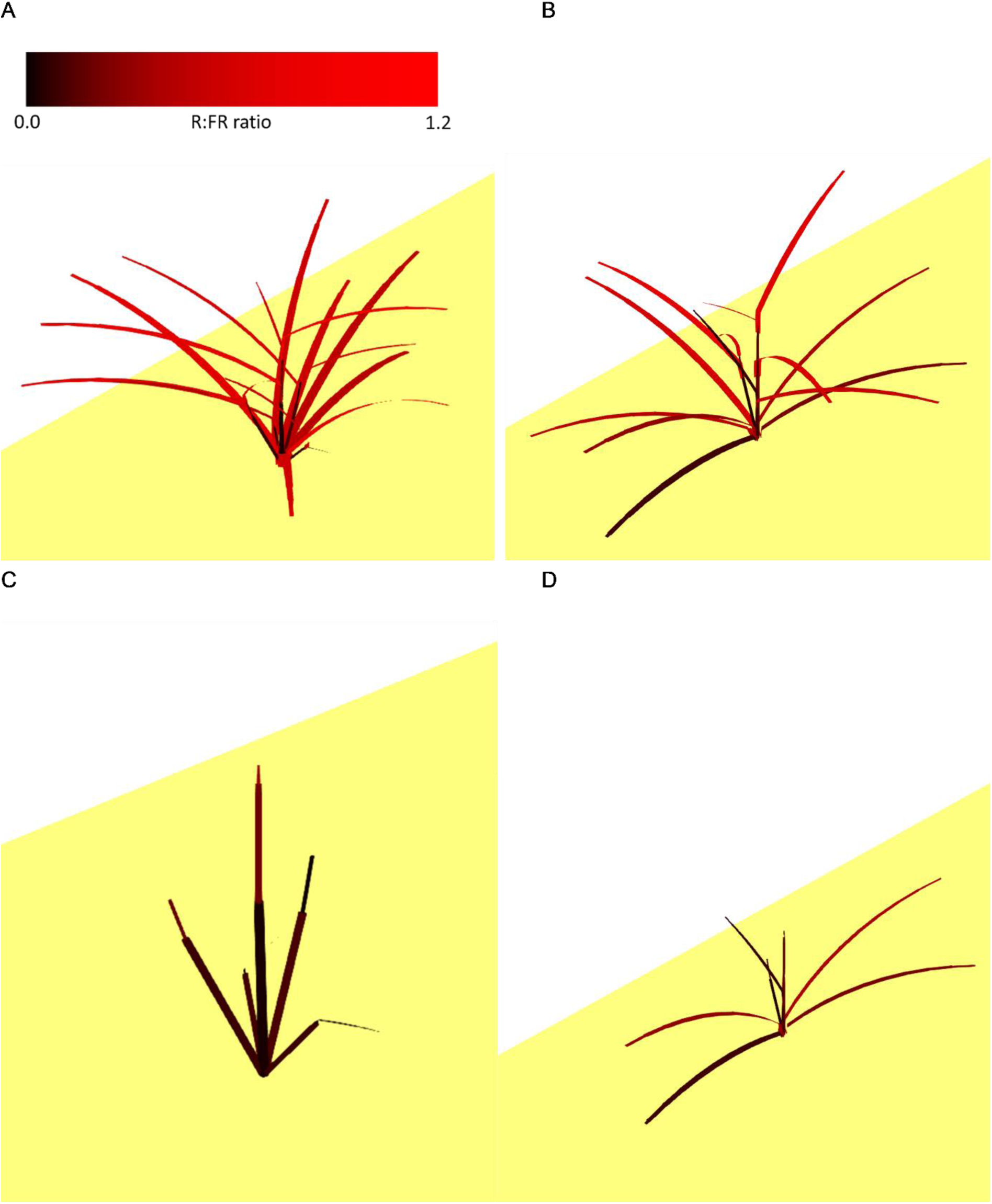
Individual cereal plants rendered from a fully simulated field at 40 DAE, colored based on organ red:far-red ratio (neighbouring plants removed for visualization purposes). A: A cereal plant from the cereal-legume treatment. B: A cereal plant from the high-density cereal sole crop treatment. C & D: The same plants as in A and B respectively, but with organs having an R:FR ratio exceeding 0.4 removed for visualization purposes.

### Competitiveness and crop performance

Simulating and analyzing two distinct species in both intercrops and sole crops, across two densities, provided insights into their competitiveness and subsequent performance. In this context, the cereal species was the stronger competitor of the two species, characterized by a higher biomass production in systems with fewer cereal plants, whether achieved through plant density or intercropping. The explicit modeling of competitive adaptation, including adaptive elongation and branching (in cereals only), resulted in diverse plant responses. A next step is to quantify the impact of specific plant traits and adaptive responses through simulations of varying cropping systems containing plants with a varying degree of competitive adaptivity. However, such an investigation extends beyond the current scope of this paper and will be presented in a forthcoming publication.

## Concluding remarks

Through this case study we can see that VPL is already mature enough to simulate intercrops using FSP modeling and it can capture many of the plant-environment interactions including photosynthesis, red:far red ratio or competition among plants for radiation. The advanced visualization system in VPL is particularly useful for gaining insights into the simulations, whereas the inclusion of a ray tracer allows for general purpose simulation of radiation interception without restrictive assumptions about the system. We are currently extending the intercrop model to account for roots and microclimate and further developing VPL in parallel to ensure that such type of modelling is also well supported.

## Supporting information

R script

## Supplement S1

**Figure S1.1.**
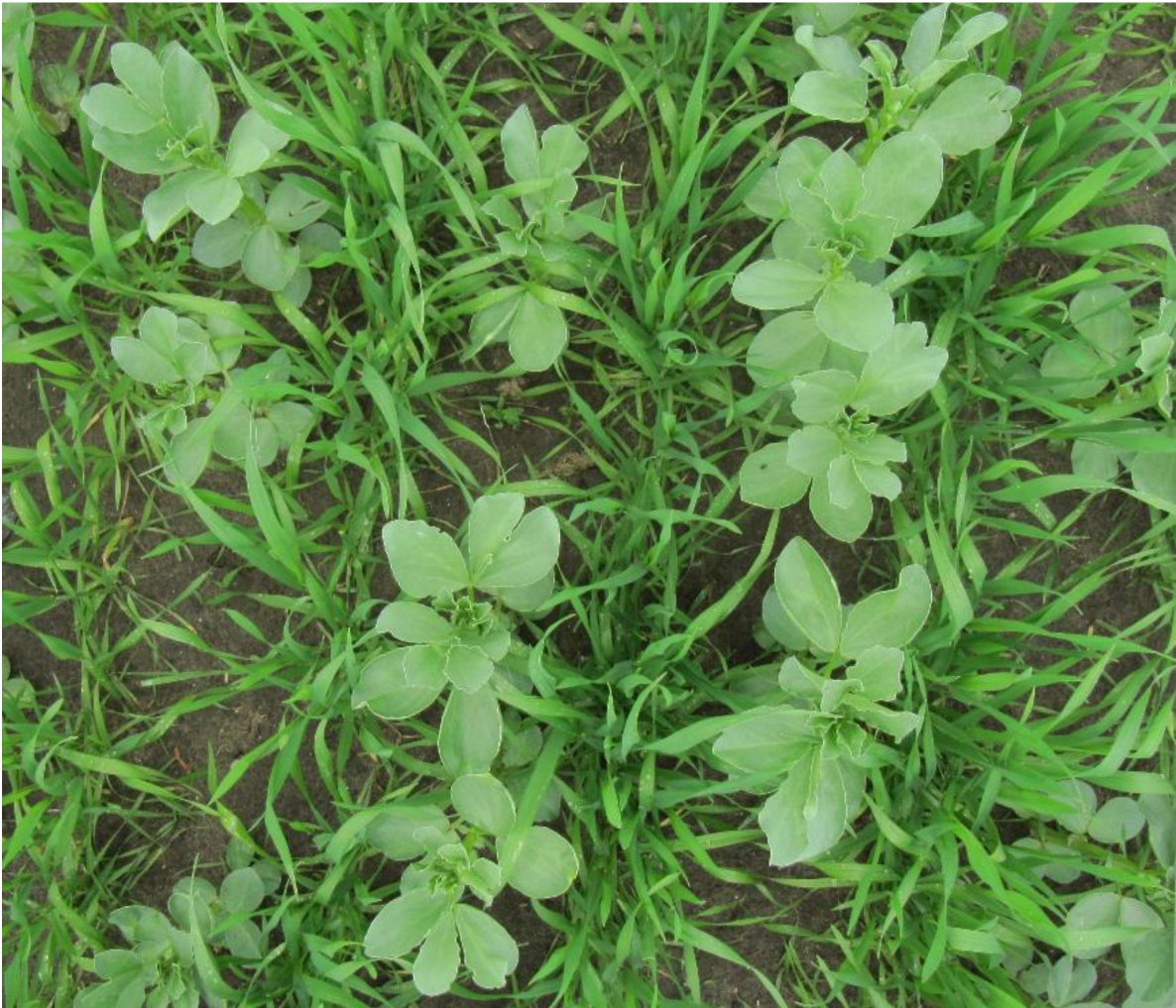
A top-down view of a triticale-faba bean row replacement intercropping system.

## Supplement 2

### Cereal-legume functional-structural plant model description

This paper presents a case study utilizing a functional-structural plant (FSP) model developed with the Virtual Plant Laboratory (VPL), in conjunction with various Julia packages (see source code for details: https://git.wur.nl/david.kottelenberg/fspm_vpl_dk). This FSP model was designed to demonstrate key features of VPL and serves as an illustrative example, and can be employed as a foundation or inspiration for new models. This version is a proof of concept and future updates are anticipated, as ongoing development focuses on refining this model and others.

This FSP model simulates at a daily time-step the growth, development, and architecture of two plant species, a cereal and a legume, with representative parameter values though no formal calibration was performed. The model describes each individual plant separately and plant-to-plant interactions occur through the radiation environment, i.e., via shading (see Growth section below for details).

At the start of a simulation, a FieldBase object is created, to store field-level variables at every time step (Fig S2.1), which is part of the model output. Each plant is represented by a Graph object and the nodes represent either geometric transformations required by the turtle graphics approach or organs (with associated geometry and internal variables). In addition, plant-level data are stored within each graph inside a PlantBase object. Information stored in PlantBase is used for plant growth, development, and visualization, as well as model output. The simulation output is comprised of the FieldBase object and an array of Graph objects (with their respective PlantBase objects), which are saved for every simulated day. This allows retrieving all the information generated during a simulation at any point, as well as restarting any simulation at any point.

Six types of organs are defined: meristem, internode, leaf, root, flower, and fruit. Organs are characterized by different variables (organized in different objects) depending on their function. All organs contain general organ information in an OrganInfo object, organs that can grow (i.e., all organs except the meristem) contain growth parameters in a GrowthVars object, and organs that can photosynthesize (i.e., internode and leaf) contain photosynthesis parameters in a PhotosynthesisVars object.

Plant development and growth is driven by air temperature, which is taken as the daily average temperature for each calendar day from the years 2001 to 2019 in De Bilt, The Netherlands (52° 06’ 36.00” N, 5° 10’ 50.02” E) (KNMI, 2024). The ensuing sections outline the processes governing plant development, growth, and architecture.

### Development

A new plant starts with a meristem representing the seed. This meristem will produce phytomers over time which represent the basic unit of structure of the aboveground part of the plant. A phytomer comprises of an internode, a leaf with an insertion angle of 50°, and a dormant lateral meristem or bud with an insertion angle of 40°. Phytomers are arranged in a spiral pattern with a rotation angle of 137.5°. Leaf appearance (phyllochron) in cereals is delayed in relation to the leaf initiation rate at the apex, and set to 90 °C d cumulative with the phytomer rank, during which they only act as a sink without active growth. Legume phyllochron is the same as the plastochron, i.e. no delay (see below).

Development of new organs is described by a graph rewriting rule that transforms meristems into new phytomers followed by this same meristem at fixed thermal time intervals (i.e., a plastochron of 45 °C d and 50 °C d for cereal and legume, respectively). The rule is as follows (we have removed the parameters of each object for clarity):

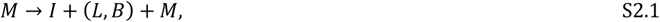

where *M* is a meristem, *I* is an internode and *L* and *B* represent a leaf and bud, respectively.

**Figure S2.1.**
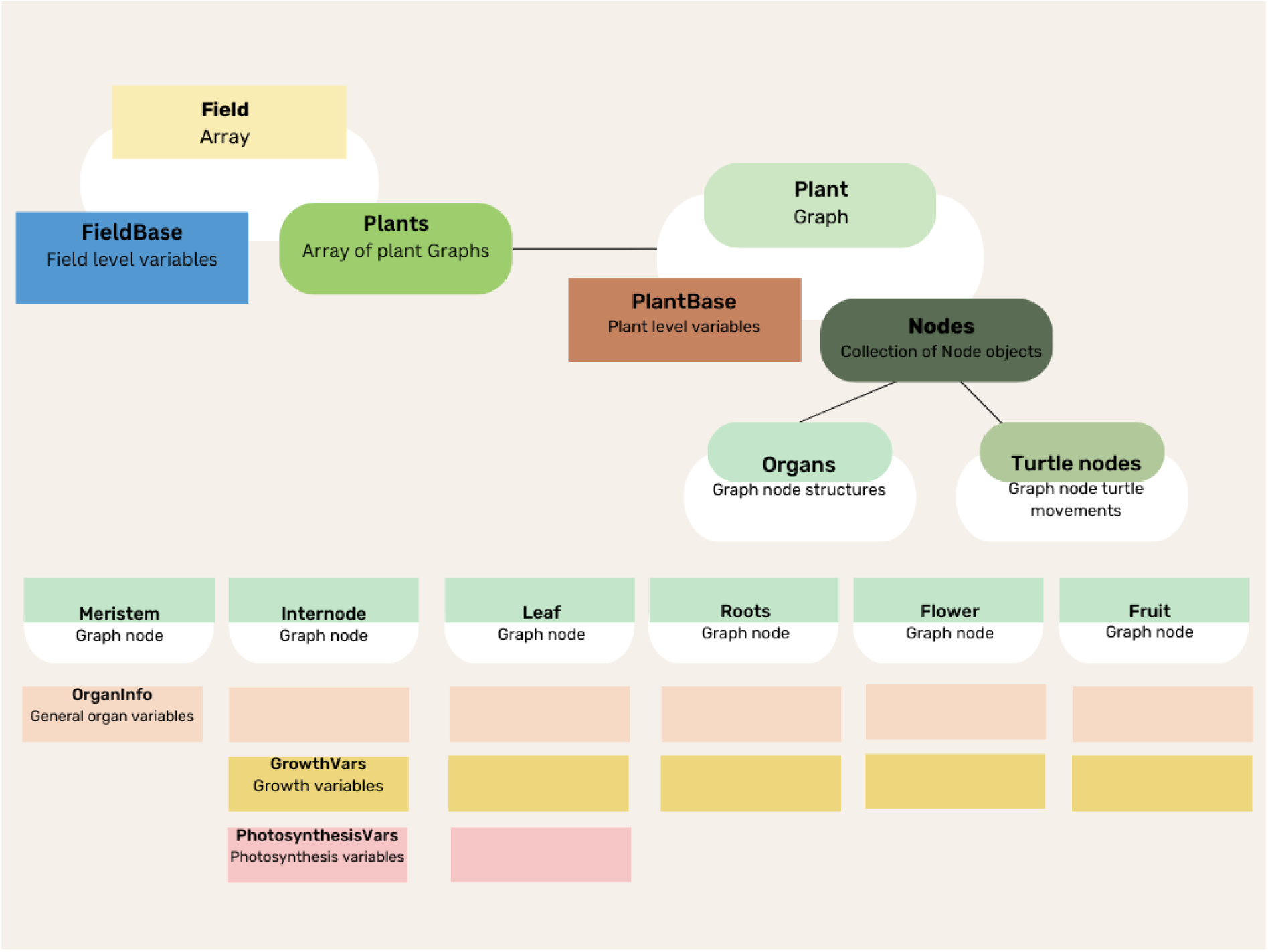
Diagram describing model structures and organization. The simulated field contains a FieldBase object and an array of plants. Each plant is represented by a Graph object containing a PlantBase object and nodes. There are two types of nodes: Turtle movement nodes and organs. Organs can be one of the following node objects: Meristem, Internode, Leaf, Roots, Flower, Fruit. The model defines various objects for organ variable storage. Organs may contain one or multiple objects, depending on their functionalities. The specified objects include: OrganInfo (storing general organ variables), GrowthVars (storing growth variables) and PhotosynthesisVars (storing photosynthesis variables).

For the cereal species, the first four phytomers of each tiller contain non-elongating internodes and they are the only phytomers that can produce tillers. Buds mature after 240 °C d and bud break is triggered in mature buds by a red:far-red ratio larger than 1.0, as detected by a 1 cm sensor around the plant base (see Growth section for details). Following the accumulation of ten phytomers for cereals and four for legumes, flowering phytomers emerge, characterized by one Flower object for cereals representing the spike, and two Flower objects per phytomer for legumes. Cereal species, being determinate, cease further phytomer development on a flowering stem. Flowers become fruits after 100 °C d (cereal) or 200 °C d (legume). Additionally, a tiller is flagged for abortion if it lacks Flower or Fruit objects and its source:sink ratio (see Growth section) drops below 1.0 (cereal) or 0.2 (legume). Branching is disabled for legumes. Finally, a plant is removed if flagged for death or upon reaching the designated harvest date, set 110 days after simulation start. A plant is flagged for death if its source:sink ratio drops below 0.01 without possessing any Fruit or Flower objects.

### Growth

The growth dynamics in this model are based on the supply (source) of and demand (sink) for assimilates, after deduction of plant maintenance respiration.

### Source

Initial assimilates for early seedling growth come from the seed endosperm, which amounts to 20 mg for cereals and 40 mg for legumes. The release of endosperm occurs over the first few days and follows a negative exponential function:

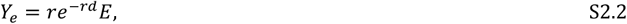

where *Y*_*e*_ (mg) represents the assimilates released from the endosperm on a given day, *r* (d^−1^) is the rate of endosperm release (assumed to be 2 d^−1^), *d* is the plant age (d), and *E* (mg) is the initial endosperm biomass.

Plants gain assimilates through photosynthesis driven by radiation capture and temperature. Radiation sources are simulated using the SkyDomes.jl package of the VPLverse. At every time step, a sky dome is created with ten million rays for diffuse radiation, assuming a standard overcast model (Moon and Spencer, 1942), and discretizing the sky into 108 sectors assuming equal solid angles (12 zenith divisions and 9 azimuth divisions). Additionally, direct radiation is modelled using a single radiation source with one million rays and this is repeated for multiple times of the day (see below for details). The calculations for radiation intensity and solar angles are based on the day of the year, the time of day, and latitude. Simulated wavebands were photosynthetic active radiation (PAR) (400-700 nm), red (600-700 nm), and far-red (701- 750 nm) all of them expressed in µmol m^−2^ s^−1^. All the calculations of solar angles, incoming radiation intensity in different wavebands and discretization of the sky hemisphere are performed by the SkyDomes.jl package.

Upon scene creation (see Architecture below), the geometries associated to Internode and Leaf objects are generated, as well as their optical properties (we assume them to be perfect Lambertian diffusers, so Lambertian objects are used). Leaves are divided into multiple sub-organs (e.g. sheath and blade), and leaf blades and leaflets are divided into multiple rectangles. In order to keep track of the absorbed radiation, each geometry element contains a distinct material object with specified transmittance and reflectance parameters (Table S2.1). For simplicity, fruits and flowers are not photosynthesizing and not capturing radiation and therefore transmit all and reflect none of the incoming radiation.

To account for diurnal variation in solar radiation and angle, total daily photosynthesis is calculated for each leaf using a Gaussian integration rule of order ten. This means that radiation absorption is computed at ten different time points throughout the daytime. For the direct radiation, the ray tracer is executed separately for each time point to account for differences in solar angle. For diffuse radiation, it is only executed once per day as the angular distribution of diffuse radiation does not change during the day (we just correct for changes in incoming diffuse radiation during the day).

At each time point of the day, assimilates generated by photosynthesis are computed for every leaf of every plant. Given that leaves are composed of multiple rectangles, photosynthesis is computed individually for each leaf part. The resulting assimilates calculated for each part are then aggregated to the entire leaf. The photosynthesis model for C3 species in the Ecophys.jl package, based on equations by Yin & Struik (2009), was used. In all calculations of photosynthesis, we assumed a relative humidity of 0.75, leaf temperature equal to the average temperature of that day, an air CO2 partial pressure of 400.0 µmol mol^−1^, oxygen of 210 mmol mol^−1^, and a boundary layer conductance to CO2 of 0.5 mol m^−2^ s^−1^.

**Table S2.1.** Transmittance and reflectance proportion parameters for the different photosynthetic organs for cereals and legumes.

| Cereal | Transmittance |  |  | Reflectance |  |  |
| --- | --- | --- | --- | --- | --- | --- |
|  | PAR | Red | Far-red | PAR | Red | Far-red |
| Internode | 0.0 | 0.0 | 0.0 | 0.10 | 0.05 | 0.80 |
| Leaf | 0.03 | 0.03 | 0.40 | 0.10 | 0.05 | 0.40 |
| Flower | 1.0 | 1.0 | 1.0 | 0.0 | 0.0 | 0.0 |
| Fruit | 1.0 | 1.0 | 1.0 | 0.0 | 0.0 | 0.0 |
| Legume |  |  |  |  |  |  |
| Internode | 0.0 | 0.0 | 0.0 | 0.15 | 0.10 | 0.80 |
| Leaf | 0.04 | 0.04 | 0.40 | 0.15 | 0.10 | 0.40 |
| Flower | 1.0 | 1.0 | 1.0 | 0.0 | 0.0 | 0.0 |
| Fruit | 1.0 | 1.0 | 1.0 | 0.0 | 0.0 | 0.0 |

### Sink

The potential and actual sink strength of organs are based on their potential growth rate (mg °C d^−1^), derived from the beta growth function as specified by Yin et al. (2003):

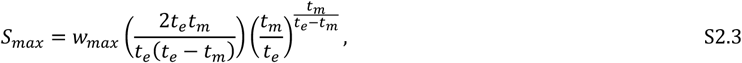

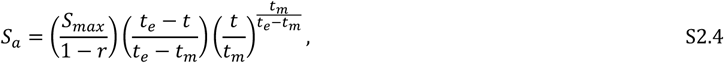

where *S*_*max*_ (mg) is the potential sink strength, *w*_*max*_ (mg) is the potential organ biomass, *t*_*e*_ (°C d) is the growth duration, *t*_*m*_ (°C d) is the age at which maximum organ growth is reached, *S*_*a*_ (mg) is realized sink strength, *r* is the proportion of biomass used for growth respiration, and *t* (°C d) is the organ age.

Plant organs are assigned potential biomass (*w*_*max*_) and growth duration (*t*_*e*_) parameters as specified in Table S2.2. Plant maintenance respiration is assumed to be 2% of the plant biomass whereas organ growth respiration is assumed to be 40% of the biomass increment across all organs. Each organ’s relative sink strength is proportional to the entire plant summed sink strength:

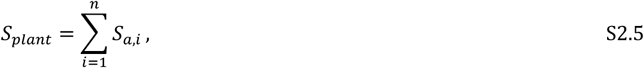

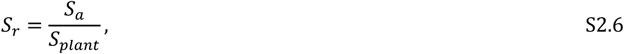

where *S*_*plant*_ (mg) is the sink strength of the plant, *n* is the total number of organs of that plant, *S*_*a,i*_ (mg) is the sink strength of the *i*-th organ, and *S*_*r*_ is the relative sink strength of an organ. An organ receives assimilates in proportion to its relative sink strength, not exceeding its absolute sink strength. Any surplus assimilates post-allocation are stored in the plant reserve pool, subsequently added to available assimilates in the next time step.

**Table S2.2.** Growth parameters for the different growing organs for cereals and legumes.

| | Potential biomass (mg) | Growth duration ($^\circ\text{C d}$) |
| --- | --- | --- |
| **Cereal** |  |  |
| Internode | 266 | 182 |
| Leaf | 266 | 220 |
| Roots | 2000 | 1800 |
| Flower | 20 | 200 |
| Fruit | 3000 | 800 |
| **Legume** |  |  |
| Internode | 1000 | 750 |
| Leaf | 1000 | 700 |
| Roots | 2000 | 1800 |
| Flower | 20 | 200 |
| Fruit | 1500 | 500 |

### Organ growth

#### Internodes

Internode elongation is derived from the increase in biomass and a fixed specific internode length. For cereals, the specific internode length is 0.6 mm mg^−1^, while for legumes it is 0.15 mm mg^−1^. Internode width is calculated as:

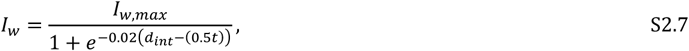

where *I*_*w*_ (m) is the internode width, *I*_*w,max*_ (m) is the maximum internode width, *d*_*int*_ (°C d) is the internode age, and *t* (°C d) is the growth duration of the internode. In this model, *I*_*w,max,cereal*_ = 0.005 m and *I*_*w,max,legume*_ = 0.015 m. Furthermore, internode elongation can respond plastically to the red:far-red ratio. To this end, the specific internode length is modulated by an elongation factor (*E*), determined by the red:far-red ratio of radiation captured by a 1 cm length sensor around the apex of the main stem. The elongation factor is defined as:

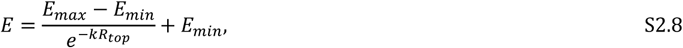

where *E*_*max*_ is the maximum elongation factor, *E*_*min*_ is the minimum elongation factor, *k* is the function shape coefficient, and *R*_*top*_ is the red:far-red ratio detected by the apex sensor. In this model, *E*_*max*_ = 2.0, *E*_*min*_ = 1.0, and *k* = 5.0.

#### Leaves

Leaf area is derived from leaf biomass and a fixed leaf mass per area set at 4.2 mg cm^−2^ (cereal) and 2.1 mg cm^−2^ (legume).

Cereals contain a leaf sheath and the fraction of the leaf biomass growth assigned to the sheath determined by the following sigmoid equation:

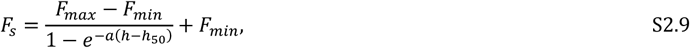

where *F*_*s*_ is the fraction of growth assigned to the leaf sheath of a particular leaf, *F*_*max*_ is the maximum fraction, *F*_*min*_ is the minimum fraction, *a* is the steepness parameter for the sigmoid curve, ℎ is the cumulative phytomer number of the leaf (i.e. the relative vertical location on the plant), and ℎ50 is the cumulative phytomer number where *F*_*s*_ is at its halfway point. This results in shorter leaf sheaths at the plant’s base, gradually increasing in size towards the top. The specified values in this model are *F*_*max*_ = 0.3, *F*_*min*_ = 0.05, *b* = 2.0, and ℎ50 = 4. Sheath length is derived from a specific sheath length of 2.5 mm mg^−2^, and their widths are set to 85% of their lengths. The length and width of cereal blades are derived from leaf area assuming a ratio between length and maximum width of 27.

In legumes, petiolules, and rachi gain assimilates as fractions of the leaf’s assigned assimilates, set at 0.001 and 0.07, respectively. Petiolules, and rachi are hollow cylinders with their lengths calculated assuming fixed specific petiolule length (1.5 mm mg^−1^) and specific rachis length (1.5 mm mg^−1^). Petiolule and rachis widths are assumed to be 1/15^th^ of their length. No petioles are assumed.

#### Roots

Roots, although not explicitly modeled, are included as sinks for assimilates (Table S2.2).

#### Flowers and fruits

Flowers serve as small sinks, indicating the initiation of flowering, and subsequently transform into fruits. Fruits, acting as substantial sinks, accumulate biomass that contributes to the eventual yield of the plant.

### Architecture

#### Internodes

Internodes are modeled as hollow cylinders, and their length and width are determined by their growth (see Growth section).

#### Leaves

The leaf blades of cereals have a narrow oblong shape, with the maximum width positioned at 72.49% of the blade length (Fig. S2.2A). Legume leaves are compound, featuring a petiole extending from the stem (assumed absent here), a rachis continuing from the petiole, petiolules perpendicular to the rachis, and leaflets at the end of each petiolule. At the end of the rachis, two leaflets are positioned at a −45 and +45 degree angle from the rachis tip (Fig. S2.2B).

The number of leaflets is based on leaf growth, with each leaf initially comprising two leaflets. When these leaflets reach their maximum size, two new leaflets are created, each with their respective petiolule. The maximum number of leaflets per leaf is eight. Leaflets are shaped as ovals, with the maximum width located at 40% of their length, and a length:width ratio of 2. Cereal leaf blades and legume leaflets have a curvature (*C*) determined by the following saturating function:

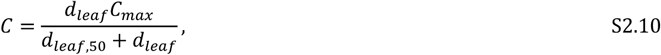

where *d*_*leaf*_ (°C d) is the leaf age, *C*_*max*_ is the maximum leaf curvature, and *d*_*leaf*,50_ is the age at which leaf curvature is half-maximum. In this model, *C*_*max,cereal*_ = 46.0, *C*_*max,legume*_ = 70.0, *d*_*leaf,50,cereal*_ = 80.0 (°C d), *d*_*leaf,50,Cereal*_ = 10.0 (°C d).

#### Flowers and Fruits

Cereal flowers resemble smaller cereal spikes, taking the form of hollow cylinders on top of the stem, featuring a cone that seals both the top and bottom of the cylinder. Legume flowers have a 45° insertion angle and are composed of by five oval shapes representing petals arranged in a circular pattern with a 60° insertion angle, creating an open flower-like structure.

The architecture of cereal fruits mirrors that of cereal flowers. The size of cereal fruits increases proportionally with biomass accumulation. Legume fruit architecture aligns with cereal fruit architecture, except with an insertion angle of 20°.

### Roots

As roots solely function as sinks in this model, no specific architecture is defined for them.

**Figure S2.2.**
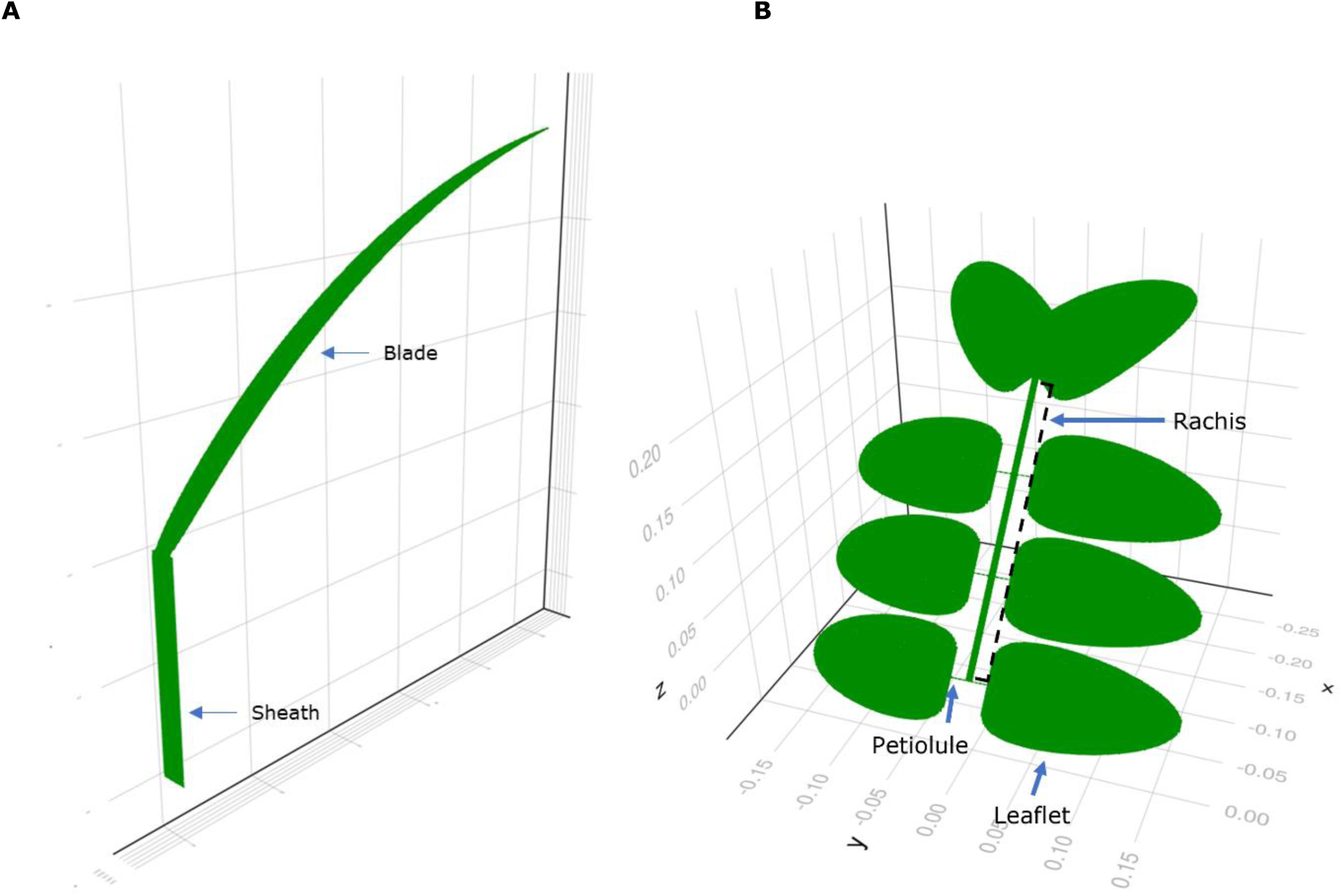
Examples of individual leaves. A: Cereal leaf, consisting of a sheath and blade. B: Legume leaf, consisting of eight leaflets which are attached to the rachis with their petiolules, except for the leaflets at the end which are directly connected to the rachis.

## Supplement S3

### Simulation replicates

Simulation replicates show similar patterns (Fig. S3.1), justifying the pooling of the data for analysis across treatments.

**Figure S3.1.**
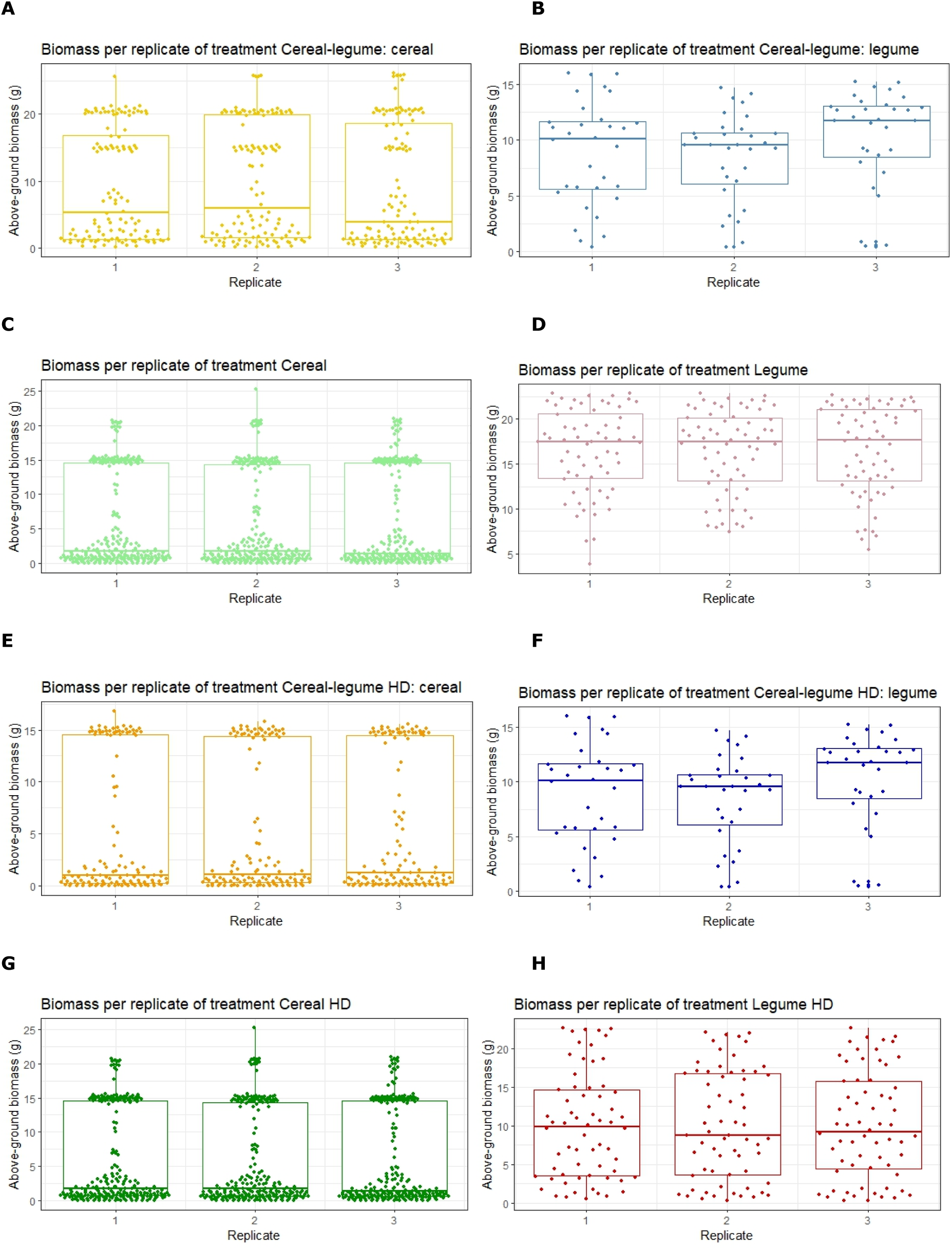
Box plots of the different treatments per replicate. A: Cereal-legume: cereal treatment. B: Cereal-legume: legume treatment. C: Cereal treatment. D: Legume treatment. E: Cereal-legume HD: cereal treatment. F: Cereal-legume HD: legume treatment. G: Cereal HD treatment. H: Legume HD treatment. HD: high density.

### Elongation dynamics

There are two aspects influencing competition in plant height. Limited light capture due to competition results in fewer resources available for elongation, as illustrated by a lower source:sink ratio of the plant. Contrastingly, environmental signals that anticipate competition e.g. through a lower red:far-red ratio as detected by the plant can induce an adaptive elongation response in the plant. Both processes were contributing to height differences between treatments in the simulations (Figures 4 and S3.2).

**Figure S3.2.**
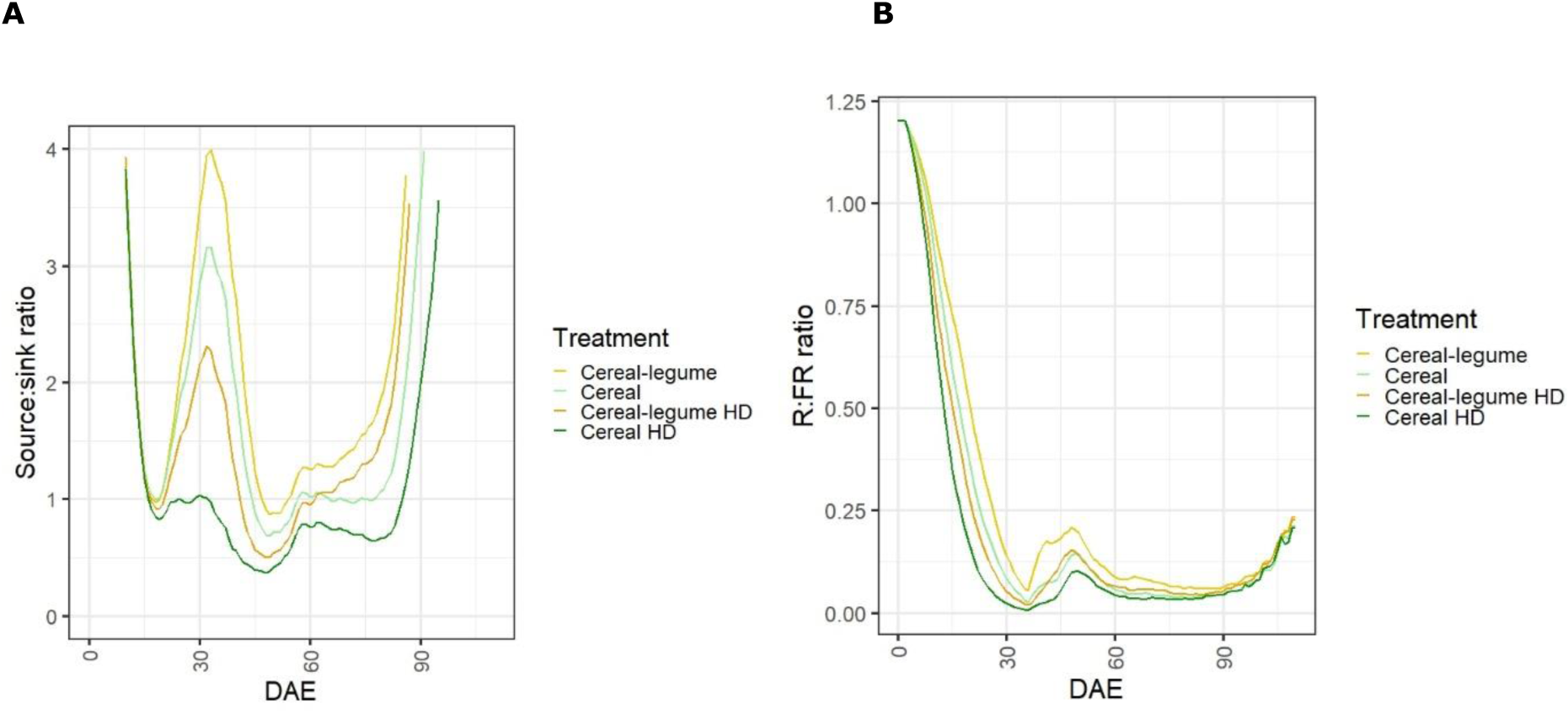
Cereal source:sink ratio and red:far-red ratio detected at the top of the plants over time in days after emergence (DAE). A: Source:sink ratio. B: Red:far-red (R:FR) ratio. HD: high density.

### LER

LER is a metric indicating how much land of monocrops is needed to produce the same amount of biomass as the intercrop. Hence LER is a measure of land use efficiency of intercrops (Li et al., 2023). LER can be computed for different outputs of interest, such as yield or biomass. LER is calculated as the sum of the relative output of the crops in the intercrop (partial LER, or pLER). For example, for biomass, and given *n* species in the intercrop:

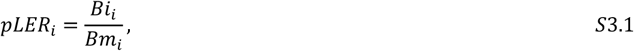

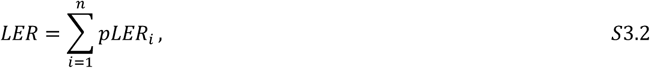

where *B*_*i*_ and *Bm*_*i*_ are the biomass of species *i* in the intercrop and monocrop setting, respectively.

