## Supplementary material for "The Virtual Plant Laboratory: a modern plant modeling framework in Julia": R script: VPL_case_study_analyses.R

```
rm(list = ls())
library(tidyverse)
library(ggrepel)
library(ggbeeswarm)
library(scales)
library(patchwork)

# Treatments
# Normal: normal density, 480 cereal plants / m2, 128 legume plants / m2
# HD: high plant density (within row plant density only), 960 cereal plants / m2, 256 legume plants / m2

# Set the location of data
dir_cereal_legume <- "C:/Users/kotte007/OneDrive - Wageningen University & Research/PhD/Git/FSPM_VPL_DK/cereal_legume_f6/"
dir_cereal <- "C:/Users/kotte007/OneDrive - Wageningen University & Research/PhD/Git/FSPM_VPL_DK/cereal_f6/"
dir_legume <- "C:/Users/kotte007/OneDrive - Wageningen University & Research/PhD/Git/FSPM_VPL_DK/legume_f6/"
dir_cereal_legume_hd <- "C:/Users/kotte007/OneDrive - Wageningen University & Research/PhD/Git/FSPM_VPL_DK/cereal_legume_hd_f6/"
dir_cereal_hd <- "C:/Users/kotte007/OneDrive - Wageningen University & Research/PhD/Git/FSPM_VPL_DK/cereal_hd_f6/"
dir_legume_hd <- "C:/Users/kotte007/OneDrive - Wageningen University & Research/PhD/Git/FSPM_VPL_DK/legume_hd_f6/"

# Read the data
## Plant
data_cereal_legume_1 <- read_delim(file = paste(dir_cereal_legume, "data_plants.txt", sep = ""), delim = "\t")
data_cereal_legume_2 <- read_delim(file = paste(dir_cereal_legume, "data_plants_1.txt", sep = ""), delim = "\t")
data_cereal_legume_3 <- read_delim(file = paste(dir_cereal_legume, "data_plants_2.txt", sep = ""), delim = "\t")
data_cereal_1 <- read_delim(file = paste(dir_cereal, "data_plants.txt", sep = ""), delim = "\t")
data_cereal_2 <- read_delim(file = paste(dir_cereal, "data_plants_1.txt", sep = ""), delim = "\t")
data_cereal_3 <- read_delim(file = paste(dir_cereal, "data_plants_2.txt", sep = ""), delim = "\t")
data_legume_1 <- read_delim(file = paste(dir_legume, "data_plants.txt", sep = ""), delim = "\t")
data_legume_2 <- read_delim(file = paste(dir_legume, "data_plants_1.txt", sep = ""), delim = "\t")
data_legume_3 <- read_delim(file = paste(dir_legume, "data_plants_2.txt", sep = ""), delim = "\t")
data_cereal_legume_hd_1 <- read_delim(file = paste(dir_cereal_legume_hd, "data_plants.txt", sep = ""), delim = "\t")
data_cereal_legume_hd_2 <- read_delim(file = paste(dir_cereal_legume_hd, "data_plants_1.txt", sep = ""), delim = "\t")
data_cereal_legume_hd_3 <- read_delim(file = paste(dir_cereal_legume_hd, "data_plants_2.txt", sep = ""), delim = "\t")
data_cereal_hd_1 <- read_delim(file = paste(dir_cereal_hd, "data_plants.txt", sep = ""), delim = "\t")
data_cereal_hd_2 <- read_delim(file = paste(dir_cereal_hd, "data_plants_1.txt", sep = ""), delim = "\t")
data_cereal_hd_3 <- read_delim(file = paste(dir_cereal_hd, "data_plants_2.txt", sep = ""), delim = "\t")
data_legume_hd_1 <- read_delim(file = paste(dir_legume_hd, "data_plants.txt", sep = ""), delim = "\t")
data_legume_hd_2 <- read_delim(file = paste(dir_legume_hd, "data_plants_1.txt", sep = ""), delim = "\t")
data_legume_hd_3 <- read_delim(file = paste(dir_legume_hd, "data_plants_2.txt", sep = ""), delim = "\t")

## Field
data_cereal_legume_field_1 <- read_delim(file = paste(dir_cereal_legume, "data_field.txt", sep = ""), delim = "\t")
data_cereal_legume_field_2 <- read_delim(file = paste(dir_cereal_legume, "data_field_1.txt", sep = ""), delim = "\t")
data_cereal_legume_field_3 <- read_delim(file = paste(dir_cereal_legume, "data_field_2.txt", sep = ""), delim = "\t")
data_cereal_field_1 <- read_delim(file = paste(dir_cereal, "data_field.txt", sep = ""), delim = "\t")
data_cereal_field_2 <- read_delim(file = paste(dir_cereal, "data_field_1.txt", sep = ""), delim = "\t")
data_cereal_field_3 <- read_delim(file = paste(dir_cereal, "data_field_2.txt", sep = ""), delim = "\t")
data_legume_field_1 <- read_delim(file = paste(dir_legume, "data_field.txt", sep = ""), delim = "\t")
data_legume_field_2 <- read_delim(file = paste(dir_legume, "data_field_1.txt", sep = ""), delim = "\t")
data_legume_field_3 <- read_delim(file = paste(dir_legume, "data_field_2.txt", sep = ""), delim = "\t")
data_cereal_legume_hd_field_1 <- read_delim(file = paste(dir_cereal_legume_hd, "data_field.txt", sep = ""), delim = "\t")
data_cereal_legume_hd_field_2 <- read_delim(file = paste(dir_cereal_legume_hd, "data_field_1.txt", sep = ""), delim = "\t")
data_cereal_legume_hd_field_3 <- read_delim(file = paste(dir_cereal_legume_hd, "data_field_2.txt", sep = ""), delim = "\t")
data_cereal_hd_field_1 <- read_delim(file = paste(dir_cereal_hd, "data_field.txt", sep = ""), delim = "\t")
data_cereal_hd_field_2 <- read_delim(file = paste(dir_cereal_hd, "data_field_1.txt", sep = ""), delim = "\t")
data_cereal_hd_field_3 <- read_delim(file = paste(dir_cereal_hd, "data_field_2.txt", sep = ""), delim = "\t")
data_legume_hd_field_1 <- read_delim(file = paste(dir_legume_hd, "data_field.txt", sep = ""), delim = "\t")
data_legume_hd_field_2 <- read_delim(file = paste(dir_legume_hd, "data_field_1.txt", sep = ""), delim = "\t")
data_legume_hd_field_3 <- read_delim(file = paste(dir_legume_hd, "data_field_2.txt", sep = ""), delim = "\t")

# Combine the simulation reps
data_cereal_legume <- bind_rows(data_cereal_legume_1,
                                data_cereal_legume_2,
                                data_cereal_legume_3
                                ) %>% 
  mutate(sim = c(rep(1, nrow(data_cereal_legume_1)),
                  rep(2, nrow(data_cereal_legume_2)),
                  rep(3, nrow(data_cereal_legume_3))
                 ))

data_cereal <- bind_rows(data_cereal_1,
                         data_cereal_2,
                         data_cereal_3) %>% 
  mutate(sim = c(rep(1, nrow(data_cereal_1)),
                  rep(2, nrow(data_cereal_2)),
                  rep(3, nrow(data_cereal_3))))

data_legume <- bind_rows(data_legume_1,
                         data_legume_2,
                         data_legume_3
                         ) %>% 
  mutate(sim = c(rep(1, nrow(data_legume_1)),
                  rep(2, nrow(data_legume_2)),
                  rep(3, nrow(data_legume_3))
                 ))

data_cereal_legume_hd <- bind_rows(data_cereal_legume_hd_1,
                                   data_cereal_legume_hd_2,
                                   data_cereal_legume_hd_3) %>% 
  mutate(sim = c(rep(1, nrow(data_cereal_legume_hd_1)),
                  rep(2, nrow(data_cereal_legume_hd_2)),
                  rep(3, nrow(data_cereal_legume_hd_3))))

data_cereal_hd <- bind_rows(data_cereal_hd_1,
                            data_cereal_hd_2,
                            data_cereal_hd_3) %>% 
  mutate(sim = c(rep(1, nrow(data_cereal_hd_1)),
                  rep(2, nrow(data_cereal_hd_2)),
                  rep(3, nrow(data_cereal_hd_3))))

data_legume_hd <- bind_rows(data_legume_hd_1,
                            data_legume_hd_2,
                            data_legume_hd_3) %>% 
  mutate(sim = c(rep(1, nrow(data_legume_hd_1)),
                  rep(2, nrow(data_legume_hd_2)),
                  rep(3, nrow(data_legume_hd_3))))

data_all <- bind_rows(data_cereal_legume,
                      data_cereal,
                      data_legume,
                      data_cereal_legume_hd,
                      data_cereal_hd,
                      data_legume_hd
                      ) %>% 
  mutate(treatment = c(rep("Cereal-legume", nrow(data_cereal_legume)),
                       rep("Cereal", nrow(data_cereal)),
                       rep("Legume", nrow(data_legume)),
                       rep("Cereal-legume HD", nrow(data_cereal_legume_hd)),
                       rep("Cereal HD", nrow(data_cereal_hd)),
                       rep("Legume HD", nrow(data_legume_hd))
                       )) %>% 
  mutate(treatment = factor(treatment, levels = c("Cereal-legume",
                                                  "Cereal",
                                                  "Legume",
                                                  "Cereal-legume HD",
                                                  "Cereal HD",
                                                  "Legume HD"
                                                  )))

## Field
data_cereal_legume_field <- bind_rows(data_cereal_legume_field_1,
                                      data_cereal_legume_field_2,
                                      data_cereal_legume_field_3
                                      ) %>% 
  mutate(sim = c(rep(1, nrow(data_cereal_legume_field_1)),
                  rep(2, nrow(data_cereal_legume_field_2)),
                  rep(3, nrow(data_cereal_legume_field_3))
                 ))

data_cereal_field <- bind_rows(data_cereal_field_1,
                               data_cereal_field_2,
                               data_cereal_field_3) %>% 
  mutate(sim = c(rep(1, nrow(data_cereal_field_1)),
                  rep(2, nrow(data_cereal_field_2)),
                  rep(3, nrow(data_cereal_field_3))))

data_legume_field <- bind_rows(data_legume_field_1,
                               data_legume_field_2,
                               data_legume_field_3
                               ) %>% 
  mutate(sim = c(rep(1, nrow(data_legume_field_1)),
                  rep(2, nrow(data_legume_field_2)),
                  rep(3, nrow(data_legume_field_3))
                 ))

data_cereal_legume_hd_field <- bind_rows(data_cereal_legume_hd_field_1,
                                         data_cereal_legume_hd_field_2,
                                         data_cereal_legume_hd_field_3) %>% 
  mutate(sim = c(rep(1, nrow(data_cereal_legume_hd_field_1)),
                  rep(2, nrow(data_cereal_legume_hd_field_2)),
                  rep(3, nrow(data_cereal_legume_hd_field_3))))

data_cereal_hd_field <- bind_rows(data_cereal_hd_field_1,
                                  data_cereal_hd_field_2,
                                  data_cereal_hd_field_3) %>% 
  mutate(sim = c(rep(1, nrow(data_cereal_hd_field_1)),
                  rep(2, nrow(data_cereal_hd_field_2)),
                  rep(3, nrow(data_cereal_hd_field_3))))

data_legume_hd_field <- bind_rows(data_legume_hd_field_1,
                                  data_legume_hd_field_2,
                                  data_legume_hd_field_3) %>% 
  mutate(sim = c(rep(1, nrow(data_legume_hd_field_1)),
                  rep(2, nrow(data_legume_hd_field_2)),
                  rep(3, nrow(data_legume_hd_field_3))))

data_all_field <- bind_rows(data_cereal_legume_field,
                            data_cereal_field,
                            data_legume_field,
                            data_cereal_legume_hd_field,
                            data_cereal_hd_field,
                            data_legume_hd_field) %>% 
  mutate(treatment = c(rep("Cereal-legume", nrow(data_cereal_legume_field)),
                       rep("Cereal", nrow(data_cereal_field)),
                       rep("Legume", nrow(data_legume_field)),
                       rep("Cereal-legume HD", nrow(data_cereal_legume_hd_field)),
                       rep("Cereal HD", nrow(data_cereal_hd_field)),
                       rep("Legume HD", nrow(data_legume_hd_field)))) %>% 
  mutate(treatment = factor(treatment, levels = c("Cereal-legume",
                                                  "Cereal",
                                                  "Legume",
                                                  "Cereal-legume HD",
                                                  "Cereal HD",
                                                  "Legume HD")))

data_yield <- data_all %>% 
  filter(yield != 0) %>% 
  mutate(above_ground_biomass = biomass - reserve_pool - root_biomass) %>% 
  mutate(yield_area = yield / 1000 / plant_area,
         above_ground_biomass_area = above_ground_biomass / 1000 / plant_area)


analyze_biomass_yield <- function(data, crop){
  data_y <- data %>% 
    dplyr::select(treatment, species_name, yield, yield_area, above_ground_biomass, above_ground_biomass_area)
  
  data_yield_crop <- data_y %>% 
    filter(species_name == crop)
  
  plot_yield_plant <- ggplot(data = data_yield_crop, aes(x = treatment, y = yield / 1000, color = treatment)) +
    geom_boxplot() +
    geom_quasirandom()
  
  plot_biomass_plant <- ggplot(data = data_yield_crop, aes(x = treatment, y = above_ground_biomass / 1000, color = treatment)) +
    geom_boxplot() +
    geom_quasirandom()
  
  plot_yield_area_plant <- ggplot(data = data_yield_crop, aes(x = treatment, y = yield_area, color = treatment)) +
    geom_boxplot() +
    geom_quasirandom()
  
  plot_biomass_area_plant <- ggplot(data = data_yield_crop, aes(x = treatment, y = above_ground_biomass_area, color = treatment)) +
    geom_boxplot() +
    geom_quasirandom()
  
  return(list(plot_yield_plant, 
              plot_yield_area_plant, 
              plot_biomass_plant, 
              plot_biomass_area_plant))
}

colors <- c("Cereal-legume: cereal" = "gold2",
            "Cereal-legume: legume" = "steelblue",
            "Cereal" = "lightgreen",
            "Legume" = "pink3",
            "Cereal-legume HD: cereal" = "orange2",
            "Cereal-legume HD: legume" = "blue3",
            "Cereal HD" = "green4",
            "Legume HD" = "red3")
colors_cereal <- c("Cereal-legume" = "gold2",
                   "Cereal" = "lightgreen",
                   "Cereal-legume HD" = "orange2",
                   "Cereal HD" = "green4")
colors_legume <- c("Cereal-legume" = "steelblue",
                   "Legume" = "pink3",
                   "Cereal-legume HD" = "blue3",
                   "Legume HD" = "red3")

cereal_plots <- analyze_biomass_yield(data_yield, "cereal")
legume_plots <- analyze_biomass_yield(data_yield, "legume")

repCLC <- ggplot(data = filter(data_yield, species_name == "cereal" & treatment == "Cereal-legume"), aes(x = sim, y = above_ground_biomass/1000, group = sim)) +
  geom_boxplot(color = colors_cereal["Cereal-legume"]) + 
  geom_quasirandom(color = colors_cereal["Cereal-legume"]) +
  theme_bw(base_size = 30) +
  labs(title = "Biomass per replicate of treatment Cereal-legume: cereal",
       y = "Above-ground biomass (g)",
       x = "Replicate",
       color = "Treatment")

repCLL <- ggplot(data = filter(data_yield, species_name == "legume" & treatment == "Cereal-legume"), aes(x = sim, y = above_ground_biomass/1000, group = sim)) +
  geom_boxplot(color = colors_legume["Cereal-legume"]) + 
  geom_quasirandom(color = colors_legume["Cereal-legume"]) +
  theme_bw(base_size = 30) +
  labs(title = "Biomass per replicate of treatment Cereal-legume: legume",
       y = "Above-ground biomass (g)",
       x = "Replicate",
       color = "Treatment")

repC <- ggplot(data = filter(data_yield, species_name == "cereal" & treatment == "Cereal"), aes(x = sim, y = above_ground_biomass/1000, group = sim)) +
  geom_boxplot(color = colors_cereal["Cereal"]) + 
  geom_quasirandom(color = colors_cereal["Cereal"]) +
  theme_bw(base_size = 30) +
  labs(title = "Biomass per replicate of treatment Cereal",
       y = "Above-ground biomass (g)",
       x = "Replicate",
       color = "Treatment")

repL <- ggplot(data = filter(data_yield, species_name == "legume" & treatment == "Legume"), aes(x = sim, y = above_ground_biomass/1000, group = sim)) +
  geom_boxplot(color = colors_legume["Legume"]) + 
  geom_quasirandom(color = colors_legume["Legume"]) +
  theme_bw(base_size = 30) +
  labs(title = "Biomass per replicate of treatment Legume",
       y = "Above-ground biomass (g)",
       x = "Replicate",
       color = "Treatment")

repCLCHD <- ggplot(data = filter(data_yield, species_name == "cereal" & treatment == "Cereal-legume HD"), aes(x = sim, y = above_ground_biomass/1000, group = sim)) +
  geom_boxplot(color = colors_cereal["Cereal-legume HD"]) + 
  geom_quasirandom(color = colors_cereal["Cereal-legume HD"]) +
  theme_bw(base_size = 30) +
  labs(title = "Biomass per replicate of treatment Cereal-legume HD: cereal",
       y = "Above-ground biomass (g)",
       x = "Replicate",
       color = "Treatment")

repCLLHD <- ggplot(data = filter(data_yield, species_name == "legume" & treatment == "Cereal-legume"), aes(x = sim, y = above_ground_biomass/1000, group = sim)) +
  geom_boxplot(color = colors_legume["Cereal-legume HD"]) + 
  geom_quasirandom(color = colors_legume["Cereal-legume HD"]) +
  theme_bw(base_size = 30) +
  labs(title = "Biomass per replicate of treatment Cereal-legume HD: legume",
       y = "Above-ground biomass (g)",
       x = "Replicate",
       color = "Treatment")

repCHD <- ggplot(data = filter(data_yield, species_name == "cereal" & treatment == "Cereal"), aes(x = sim, y = above_ground_biomass/1000, group = sim)) +
  geom_boxplot(color = colors_cereal["Cereal HD"]) + 
  geom_quasirandom(color = colors_cereal["Cereal HD"]) +
  theme_bw(base_size = 30) +
  labs(title = "Biomass per replicate of treatment Cereal HD",
       y = "Above-ground biomass (g)",
       x = "Replicate",
       color = "Treatment")

repLHD <- ggplot(data = filter(data_yield, species_name == "legume" & treatment == "Legume HD"), aes(x = sim, y = above_ground_biomass/1000, group = sim)) +
  geom_boxplot(color = colors_legume["Legume HD"]) + 
  geom_quasirandom(color = colors_legume["Legume HD"]) +
  theme_bw(base_size = 30) +
  labs(title = "Biomass per replicate of treatment Legume HD",
       y = "Above-ground biomass (g)",
       x = "Replicate",
       color = "Treatment")


(cereal_yield_plant <- cereal_plots[[1]] +
    theme_bw(base_size = 30) +
    theme(axis.text.x = element_text(angle = 90, vjust = 0.5, hjust=1)) +
    labs(title = "",
         y = "Grain biomass (g)",
         x = "",
         color = "Treatment") +
    scale_color_manual(values = c(colors_cereal["Cereal-legume"], 
                                  colors_cereal["Cereal"],
                                  colors_cereal["Cereal-legume HD"],
                                  colors_cereal["Cereal HD"])))

(cereal_yield_area_plant <- cereal_plots[[2]] +
    theme_bw(base_size = 30) +
    theme(axis.text.x = element_text(angle = 90, vjust = 0.5, hjust=1)) +
    labs(title = "",
         y = bquote("Grain biomass (g  "~m^-2~")"),
         x = "",
         color = "Treatment") +
    scale_color_manual(values = c(colors_cereal["Cereal-legume"], 
                                  colors_cereal["Cereal"],
                                  colors_cereal["Cereal-legume HD"],
                                  colors_cereal["Cereal HD"])))

(cereal_biomass_plant <- cereal_plots[[3]] +
    theme_bw(base_size = 30) +
    theme(axis.text.x = element_text(angle = 90, vjust = 0.5, hjust=1)) +
    labs(title = "",
         y = "Above-ground biomass (g)",
         x = "",
         color = "Treatment") +
    scale_color_manual(values = c(colors_cereal["Cereal-legume"], 
                                  colors_cereal["Cereal"],
                                  colors_cereal["Cereal-legume HD"],
                                  colors_cereal["Cereal HD"])))

(cereal_biomass_area_plant <- cereal_plots[[4]] +
    theme_bw(base_size = 30) +
    theme(axis.text.x = element_text(angle = 90, vjust = 0.5, hjust=1)) +
    labs(title = "",
         y = bquote("Above-ground biomass (g  "~m^-2~")"),
         x = "",
         color = "Treatment") +
    scale_color_manual(values = c(colors_cereal["Cereal-legume"], 
                                  colors_cereal["Cereal"],
                                  colors_cereal["Cereal-legume HD"],
                                  colors_cereal["Cereal HD"])))

(legume_yield_plant <- legume_plots[[1]] +
    theme_bw(base_size = 30) +
    theme(axis.text.x = element_text(angle = 90, vjust = 0.5, hjust=1)) +
    labs(title = "",
         y = "Grain biomass (g)",
         x = "",
         color = "Treatment") +
    scale_color_manual(values = c(colors_legume["Cereal-legume"], 
                                  colors_legume["Legume"],
                                  colors_legume["Cereal-legume HD"],
                                  colors_legume["Legume HD"])))

(legume_yield_area_plant <- legume_plots[[2]] +
    theme_bw(base_size = 30) +
    theme(axis.text.x = element_text(angle = 90, vjust = 0.5, hjust=1)) +
    labs(title = "",
         y = bquote("Grain biomass (g  "~m^-2~")"),
         x = "",
         color = "Treatment") +
    scale_color_manual(values = c(colors_legume["Cereal-legume"], 
                                  colors_legume["Legume"],
                                  colors_legume["Cereal-legume HD"],
                                  colors_legume["Legume HD"])))

(legume_biomass_plant <- legume_plots[[3]] +
    theme_bw(base_size = 30) +
    theme(axis.text.x = element_text(angle = 90, vjust = 0.5, hjust=1)) +
    labs(title = "",
         y = "Above-ground biomass (g)",
         x = "",
         color = "Treatment") +
    scale_color_manual(values = c(colors_legume["Cereal-legume"], 
                                  colors_legume["Legume"],
                                  colors_legume["Cereal-legume HD"],
                                  colors_legume["Legume HD"])))

(legume_biomass_area_plant <- legume_plots[[4]] +
    theme_bw(base_size = 30) +
    theme(axis.text.x = element_text(angle = 90, vjust = 0.5, hjust=1)) +
    labs(title = "",
         y = bquote("Above-ground biomass (g  "~m^-2~")"),
         x = "",
         color = "Treatment") +
    scale_color_manual(values = c(colors_legume["Cereal-legume"], 
                                  colors_legume["Legume"],
                                  colors_legume["Cereal-legume HD"],
                                  colors_legume["Legume HD"])))

## Histograms
data_yield_t <- data_yield %>% 
  dplyr::select(treatment, species_name, yield, yield_area, above_ground_biomass) %>% 
  mutate(treatment = ifelse(treatment %in% c("Cereal-legume", "Cereal-legume HD"), paste(treatment, species_name, sep = ": "), paste(treatment))) %>% 
  mutate(treatment = factor(treatment, levels = c("Cereal-legume: cereal",
                                                  "Cereal",
                                                  "Cereal-legume HD: cereal",
                                                  "Cereal HD",
                                                  "Cereal-legume: legume",
                                                  "Legume",
                                                  "Cereal-legume HD: legume",
                                                  "Legume HD")))

(biomass_hist <- ggplot(data = data_yield_t, aes(x = above_ground_biomass / 1000, fill = treatment)) +
    geom_histogram() +
    facet_wrap(~ treatment, nrow = 4) +
    theme_bw(base_size = 30) +
    labs(title = "",
         y = "Frequency",
         x = "Above-ground biomass (g)",
         fill = "Treatment") +
    scale_color_manual(values = c(colors["Cereal-legume: cereal"], 
                                  colors["Cereal"],
                                  colors["Cereal-legume HD: cereal"],
                                  colors["Cereal HD"],
                                  colors["Cereal-legume: legume"], 
                                  colors["Legume"],
                                  colors["Cereal-legume HD: legume"],
                                  colors["Legume HD"])))

# Field biomass
field_species_names <- sapply(1:nrow(data_all_field), function(i){
  tr <- data_all_field$treatment[i]
  sp <- data_all_field$species[i]
  if(tr %in% c("Cereal-legume", "Cereal-legume HD")){
    new_species <- ifelse(sp == 1, "cereal", "legume")
  }else{
    new_species = tolower(gsub(" HD", "", tr))
  }
  return(new_species)
})
field_treatment_names <- sapply(1:nrow(data_all_field), function(i){
  tr <- data_all_field$treatment[i]
  sp <- data_all_field$species[i]
  if(tr %in% c("Cereal-legume", "Cereal-legume HD")){
    new_treatment <- paste(tr, ifelse(sp == 1, "cereal", "legume"), sep = ": ")
  }else{
    new_treatment = paste(tr)
  }
  return(new_treatment)
})

data_yield_field <- data_all_field %>% 
  mutate(species_name = field_species_names,
         treatment = field_treatment_names) %>% 
  filter(yield != 0) %>% 
  dplyr::select(time, treatment, species_name, sim, area, yield) %>% 
  mutate(yield_area = yield / area)

corrected_yield <- sapply(1:nrow(data_yield_field), function(i){
  tr <- data_yield_field$treatment[i]
  yld <- data_yield_field$yield_area[i]
  if(grepl("-", tr)){
    new_yield <- yld * 2
  }else{
    new_yield <- yld
  }
  return(new_yield)
})

data_yield_field_corr <- data_yield_field %>% 
  mutate(yield_area = corrected_yield)

data_yield_field_agg <- aggregate(yield_area ~ treatment + species_name, data = data_yield_field_corr, FUN = mean) %>% 
  mutate(SD = aggregate(yield_area ~ treatment + species_name, data = data_yield_field_corr, FUN = sd)$yield) %>% 
  mutate(treatment = factor(treatment, levels = c("Cereal-legume: cereal",
                                                  "Cereal",
                                                  "Cereal-legume HD: cereal",
                                                  "Cereal HD",
                                                  "Cereal-legume: legume",
                                                  "Legume",
                                                  "Cereal-legume HD: legume",
                                                  "Legume HD")))

(plot_yield_field_cereal <- ggplot(data = data_yield_field_agg, aes(x = treatment, y = yield_area, fill = treatment)) +
  geom_bar(stat = "identity") +
  geom_errorbar(aes(x = treatment, ymin = yield_area - SD, ymax = yield_area + SD), width = 0.2) +
  theme_bw(base_size = 30) +
  theme(axis.text.x = element_text(angle = 90, vjust = 0.5, hjust=1)) +
  labs(x = "",
       y = bquote("Grain biomass (g  "~m^-2~")"),
       fill = "Treatment") +
    scale_fill_manual(values = c(colors["Cereal-legume: cereal"], 
                                  colors["Cereal"],
                                  colors["Cereal-legume HD: cereal"],
                                  colors["Cereal HD"],
                                  colors["Cereal-legume: legume"], 
                                  colors["Legume"],
                                  colors["Cereal-legume HD: legume"],
                                  colors["Legume HD"])))

# LAI
data_LAI <- data_all %>% 
  dplyr::select(treatment, time, species_name, leaf_area, plant_area) %>% 
  mutate(LAI = leaf_area / plant_area)

data_LAI_agg <- aggregate(LAI ~ time + species_name + treatment, data = data_LAI, FUN = mean)

(cereal_LAI <- ggplot(data = filter(data_LAI_agg, species_name == "cereal"), aes(x = time, y = LAI, color = treatment)) +
    geom_line(linewidth = 1) +
    theme_bw(base_size = 30) +
    theme(axis.text.x = element_text(angle = 90, vjust = 0.5, hjust=1)) +
    labs(title = "",
         y = "LAI",
         x = "DAE",
         color = "Treatment") +
    scale_color_manual(values = c(colors_cereal["Cereal-legume"], 
                                  colors_cereal["Cereal"],
                                  colors_cereal["Cereal-legume HD"],
                                  colors_cereal["Cereal HD"])))

(legume_LAI <- ggplot(data = filter(data_LAI_agg, species_name == "legume"), aes(x = time, y = LAI, color = treatment)) +
    geom_line(linewidth = 1) +
    theme_bw(base_size = 30) +
    theme(axis.text.x = element_text(angle = 90, vjust = 0.5, hjust=1)) +
    labs(title = "",
         y = "LAI",
         x = "DAE",
         color = "Treatment") +
    scale_color_manual(values = c(colors_legume["Cereal-legume"], 
                                  colors_legume["Legume"],
                                  colors_legume["Cereal-legume HD"],
                                  colors_legume["Legume HD"])))

new_treatments <- sapply(1:nrow(data_LAI_agg), function(i){
  tr <- data_LAI_agg$treatment[i]
  sp <- data_LAI_agg$species_name[i]
  if(grepl("-", tr)){
    return(paste(tr, sp, sep = ": "))
  }else{
    return(paste(tr, sep = ""))
  }
})
data_LAI_agg_comb <- data_LAI_agg %>% 
  mutate(treatment = new_treatments) %>% 
  mutate(treatment = factor(treatment, levels = c("Cereal-legume: cereal",
                                                  "Cereal-legume: legume",
                                                  "Cereal",
                                                  "Legume",
                                                  "Cereal-legume HD: cereal",
                                                  "Cereal-legume HD: legume",
                                                  "Cereal HD",
                                                  "Legume HD")))

(combined_LAI <- ggplot(data = data_LAI_agg_comb, aes(x = time, y = LAI, color = treatment, linetype = species_name)) +
    geom_line(linewidth = 1) +
    theme_bw(base_size = 30) +
    theme(axis.text.x = element_text(angle = 90, vjust = 0.5, hjust=1)) +
    labs(title = "",
         y = "LAI",
         x = "DAE",
         color = "Treatment",
         linetype = "Crop") +
    scale_linetype_manual(values = c(1, 2), labels = c("Cereal", "Legume")) +
    scale_color_manual(values = c(colors["Cereal-legume: cereal"], 
                                  colors["Cereal"],
                                  colors["Cereal-legume HD: cereal"],
                                  colors["Cereal HD"],
                                  colors["Cereal-legume: legume"], 
                                  colors["Legume"],
                                  colors["Cereal-legume HD: legume"],
                                  colors["Legume HD"])))

# Tillering
data_tillering <- data_all %>% 
  dplyr::select(treatment, time, species_name, nbranches, bottom_rfr_ratio)

data_tillering_agg <- aggregate(nbranches ~ time + species_name + treatment, data = data_tillering, FUN = mean)
data_rfr_agg <- aggregate(bottom_rfr_ratio ~ time + species_name + treatment, data = data_tillering, FUN = mean)

(cereal_tillering <- ggplot(data = filter(data_tillering_agg, species_name == "cereal"), aes(x = time, y = nbranches, color = treatment)) +
    geom_line(linewidth = 1) +
    theme_bw(base_size = 30) +
    theme(axis.text.x = element_text(angle = 90, vjust = 0.5, hjust=1)) +
    labs(title = "",
         y = "N tillers",
         x = "DAE",
         color = "Treatment") +
    scale_color_manual(values = c(colors_cereal["Cereal-legume"], 
                                  colors_cereal["Cereal"],
                                  colors_cereal["Cereal-legume HD"],
                                  colors_cereal["Cereal HD"])))

data_tillering <- data_all %>% 
  dplyr::select(treatment, time, species_name, nbranches, bottom_rfr_ratio)

data_tillering_agg <- aggregate(nbranches ~ time + species_name + treatment, data = data_tillering, FUN = mean)
data_rfr_agg <- aggregate(bottom_rfr_ratio ~ time + species_name + treatment, data = data_tillering, FUN = mean)

(cereal_tillering <- ggplot(data = filter(data_tillering_agg, species_name == "cereal"), aes(x = time, y = nbranches, color = treatment)) +
    geom_line(linewidth = 1) +
    theme_bw(base_size = 30) +
    theme(axis.text.x = element_text(angle = 90, vjust = 0.5, hjust=1)) +
    labs(title = "",
         y = "N tillers",
         x = "DAE",
         color = "Treatment") +
    scale_color_manual(values = c(colors_cereal["Cereal-legume"], 
                                  colors_cereal["Cereal"],
                                  colors_cereal["Cereal-legume HD"],
                                  colors_cereal["Cereal HD"])))
(cereal_rfr <- ggplot(data = filter(data_rfr_agg, species_name == "cereal"), aes(x = time, y = bottom_rfr_ratio, color = treatment)) +
    geom_line(linewidth = 1) +
    theme_bw(base_size = 30) +
    theme(axis.text.x = element_text(angle = 90, vjust = 0.5, hjust=1)) +
    labs(title = "",
         y = "R:FR ratio",
         x = "DAE",
         color = "Treatment") +
    scale_color_manual(values = c(colors_cereal["Cereal-legume"], 
                                  colors_cereal["Cereal"],
                                  colors_cereal["Cereal-legume HD"],
                                  colors_cereal["Cereal HD"])))

data_tillering_sim <- data_all %>% 
  dplyr::select(treatment, time, species_name, nbranches, bottom_rfr_ratio, sim)

data_tillering_comb <- aggregate(nbranches ~ treatment + sim + time + species_name, data = data_tillering_sim, FUN = mean) %>% 
  filter(time == 60 & species_name == "cereal")
  
data_tillering_comb_avg <- aggregate(nbranches ~ treatment, data = data_tillering_comb, FUN = mean) %>%
  mutate(SD = aggregate(nbranches ~ treatment, data = data_tillering_comb, FUN = sd)$nbranches)

(cereal_tillering <- ggplot(data = data_tillering_comb_avg, aes(x = treatment, y = nbranches, fill = treatment)) +
    geom_bar(stat = "identity") +
    geom_errorbar(aes(x = treatment, ymin = nbranches - SD, ymax = nbranches + SD), col = "black", width = 0.2) +
    theme_bw(base_size = 30) +
    theme(axis.text.x = element_text(angle = 90, vjust = 0.5, hjust=1)) +
    labs(title = "",
         y = "N tillers",
         x = "",
         fill = "Treatment") +
    scale_fill_manual(values = c(colors_cereal["Cereal-legume"], 
                                  colors_cereal["Cereal"],
                                  colors_cereal["Cereal-legume HD"],
                                  colors_cereal["Cereal HD"])))

# Height
data_height <- data_all %>% 
  dplyr::select(treatment, time, species_name, height, top_rfr_ratio, elongation_plasticity_coef, source_sink_ratio)

data_height_agg <- aggregate(height ~ time + species_name + treatment, data = data_height, FUN = mean)
data_top_rfr_agg <- aggregate(top_rfr_ratio ~ time + species_name + treatment, data = data_height, FUN = mean)
data_coef_agg <- aggregate(elongation_plasticity_coef ~ time + species_name + treatment, data = data_height, FUN = mean)
data_ssr_agg <- aggregate(source_sink_ratio ~ time + species_name + treatment, data = data_height, FUN = mean)

(cereal_height <- ggplot(data = filter(data_height_agg, species_name == "cereal"), aes(x = time, y = height, color = treatment)) +
    geom_line(linewidth = 1) +
    theme_bw(base_size = 30) +
    theme(axis.text.x = element_text(angle = 90, vjust = 0.5, hjust=1)) +
    labs(title = "",
         y = "Height (m)",
         x = "DAE",
         color = "Treatment") +
    scale_color_manual(values = c(colors_cereal["Cereal-legume"], 
                                  colors_cereal["Cereal"],
                                  colors_cereal["Cereal-legume HD"],
                                  colors_cereal["Cereal HD"])))

(cereal_top_rfr <- ggplot(data = filter(data_top_rfr_agg, species_name == "cereal"), aes(x = time, y = top_rfr_ratio, color = treatment)) +
    geom_line(linewidth = 1) +
    theme_bw(base_size = 30) +
    theme(axis.text.x = element_text(angle = 90, vjust = 0.5, hjust=1)) +
    labs(title = "",
         y = "R:FR ratio",
         x = "DAE",
         color = "Treatment") +
    scale_color_manual(values = c(colors_cereal["Cereal-legume"], 
                                  colors_cereal["Cereal"],
                                  colors_cereal["Cereal-legume HD"],
                                  colors_cereal["Cereal HD"])))

(cereal_coef <- ggplot(data = filter(data_coef_agg, species_name == "cereal"), aes(x = time, y = elongation_plasticity_coef, color = treatment)) +
    geom_line(linewidth = 1) +
    theme_bw(base_size = 30) +
    theme(axis.text.x = element_text(angle = 90, vjust = 0.5, hjust=1)) +
    labs(title = "",
         y = "Elongation coefficient",
         x = "DAE",
         color = "Treatment") +
    scale_color_manual(values = c(colors_cereal["Cereal-legume"], 
                                  colors_cereal["Cereal"],
                                  colors_cereal["Cereal-legume HD"],
                                  colors_cereal["Cereal HD"])))

(cereal_ssr <- ggplot(data = filter(data_ssr_agg, species_name == "cereal"), aes(x = time, y = source_sink_ratio, color = treatment)) +
    geom_line(linewidth = 1) +
    theme_bw(base_size = 30) +
    theme(axis.text.x = element_text(angle = 90, vjust = 0.5, hjust=1)) +
    labs(title = "",
         y = "Source:sink ratio",
         x = "DAE",
         color = "Treatment") +
    ylim(c(0, 4)) +
    scale_color_manual(values = c(colors_cereal["Cereal-legume"], 
                                  colors_cereal["Cereal"],
                                  colors_cereal["Cereal-legume HD"],
                                  colors_cereal["Cereal HD"])))


(legume_height <- ggplot(data = filter(data_height_agg, species_name == "legume"), aes(x = time, y = height, color = treatment)) +
    geom_line(linewidth = 1) +
    theme_bw(base_size = 30) +
    theme(axis.text.x = element_text(angle = 90, vjust = 0.5, hjust=1)) +
    labs(title = "",
         y = "Height (m)",
         x = "DAE",
         color = "Treatment") +
    scale_color_manual(values = c(colors_legume["Cereal-legume"], 
                                  colors_legume["Legume"],
                                  colors_legume["Cereal-legume HD"],
                                  colors_legume["Legume HD"])))

(legume_top_rfr <- ggplot(data = filter(data_top_rfr_agg, species_name == "legume"), aes(x = time, y = top_rfr_ratio, color = treatment)) +
    geom_line(linewidth = 1) +
    theme_bw(base_size = 30) +
    theme(axis.text.x = element_text(angle = 90, vjust = 0.5, hjust=1)) +
    labs(title = "",
         y = "R:FR ratio",
         x = "DAE",
         color = "Treatment") +
    scale_color_manual(values = c(colors_legume["Cereal-legume"], 
                                  colors_legume["Legume"],
                                  colors_legume["Cereal-legume HD"],
                                  colors_legume["Legume HD"])))

(legume_coef <- ggplot(data = filter(data_coef_agg, species_name == "legume"), aes(x = time, y = elongation_plasticity_coef, color = treatment)) +
    geom_line(linewidth = 1) +
    theme_bw(base_size = 30) +
    theme(axis.text.x = element_text(angle = 90, vjust = 0.5, hjust=1)) +
    labs(title = "",
         y = "Elongation coefficient",
         x = "DAE",
         color = "Treatment") +
    scale_color_manual(values = c(colors_legume["Cereal-legume"], 
                                  colors_legume["Legume"],
                                  colors_legume["Cereal-legume HD"],
                                  colors_legume["Legume HD"])))

(legume_ssr <- ggplot(data = filter(data_ssr_agg, species_name == "legume"), aes(x = time, y = source_sink_ratio, color = treatment)) +
    geom_line(linewidth = 1) +
    theme_bw(base_size = 30) +
    theme(axis.text.x = element_text(angle = 90, vjust = 0.5, hjust=1)) +
    labs(title = "",
         y = "Source:sink ratio",
         x = "DAE",
         color = "Treatment") +
    scale_color_manual(values = c(colors_legume["Cereal-legume"], 
                                  colors_legume["Legume"],
                                  colors_legume["Cereal-legume HD"],
                                  colors_legume["Legume HD"])) +
  ylim(c(0, 10)))

## LER
pLERCereal <- filter(data_cereal_legume_field, time == 110 & species == 1)$above_ground_biomass /
  filter(data_cereal_field, time == 110)$above_ground_biomass
pLERLegume <- 
  filter(data_cereal_legume_field, time == 110 & species == 2)$above_ground_biomass /
  filter(data_legume_field, time == 110)$above_ground_biomass
LER <- pLERCereal + pLERLegume

pLERCerealHD <- filter(data_cereal_legume_hd_field, time == 110 & species == 1)$above_ground_biomass /
  filter(data_cereal_hd_field, time == 110)$above_ground_biomass
pLERLegumeHD <- 
  filter(data_cereal_legume_hd_field, time == 110 & species == 2)$above_ground_biomass /
  filter(data_legume_hd_field, time == 110)$above_ground_biomass
LERHD <- pLERCerealHD + pLERLegumeHD

dataLER <- tibble(Crop = c("Cereal", "Legume", "Cereal HD", "Legume HD"),
                  Treatment = c("Cereal-legume", "Cereal-legume", "Cereal-legume HD", "Cereal-legume HD"),
                  LER = c(mean(pLERCereal), mean(pLERLegume), mean(pLERCerealHD), mean(pLERLegumeHD)),
                  SD = c(sd(pLERCereal), sd(pLERLegume), sd(pLERCerealHD), sd(pLERLegumeHD)))
                  
(LERs <- ggplot(data = dataLER, aes(x = Treatment, y = LER, fill = Crop)) + 
    geom_bar(stat = "identity") +
    geom_errorbar(aes(ymin = c(rep(dataLER$LER[1] + dataLER$LER[2], 2), rep(dataLER$LER[3] + dataLER$LER[4], 2)) - c(rep(sd(pLERCereal + pLERLegume), 2), rep(sd(pLERCerealHD + pLERLegumeHD), 2)), ymax = c(rep(dataLER$LER[1] + dataLER$LER[2], 2), rep(dataLER$LER[3] + dataLER$LER[4], 2)) + c(rep(sd(pLERCereal + pLERLegume), 2), rep(sd(pLERCerealHD + pLERLegumeHD), 2))), width = 0.2) +
    scale_fill_manual(values = c(colors["Legume"], 
                                  colors["Cereal"],
                                  colors["Legume HD"],
                                  colors["Cereal HD"])) +
    labs(x = "",
         y = "(p)LER") +
    theme_bw(base_size = 30) +
    geom_hline(yintercept = 1.0, linetype = "dashed"))


# Create figures
setwd("C:/Users/kotte007/OneDrive - Wageningen University & Research/PhD/2023/Papers/VPL")

jpeg("figures/cereal_yield_plant.jpg", units = "px", width = 900, height = 750, quality = 300)
cereal_yield_plant
dev.off()

jpeg("figures/cereal_biomass_plant.jpg", units = "px", width = 900, height = 750, quality = 300)
cereal_biomass_plant
dev.off()

jpeg("figures/cereal_yield_area_plant.jpg", units = "px", width = 900, height = 750, quality = 300)
cereal_yield_area_plant
dev.off()

jpeg("figures/cereal_biomass_area_plant.jpg", units = "px", width = 900, height = 750, quality = 300)
cereal_biomass_area_plant
dev.off()


jpeg("figures/legume_yield_plant.jpg", units = "px", width = 900, height = 750, quality = 300)
legume_yield_plant
dev.off()

jpeg("figures/legume_biomass_plant.jpg", units = "px", width = 900, height = 750, quality = 300)
legume_biomass_plant
dev.off()

jpeg("figures/legume_yield_area_plant.jpg", units = "px", width = 900, height = 750, quality = 300)
legume_yield_area_plant
dev.off()

jpeg("figures/legume_biomass_area_plant.jpg", units = "px", width = 900, height = 750, quality = 300)
legume_biomass_area_plant
dev.off()


jpeg("figures/plot_yield_field_cereal.jpg", units = "px", width = 900, height = 750, quality = 300)
plot_yield_field_cereal
dev.off()

jpeg("figures/biomass_hist.jpg", units = "px", width = 900, height = 750, quality = 300)
biomass_hist
dev.off()

jpeg("figures/cereal_LAI.jpg", units = "px", width = 900, height = 750, quality = 300)
cereal_LAI
dev.off()

jpeg("figures/legume_LAI.jpg", units = "px", width = 900, height = 750, quality = 300)
legume_LAI
dev.off()

jpeg("figures/combined_LAI.jpg", units = "px", width = 900, height = 750, quality = 300)
combined_LAI
dev.off()

jpeg("figures/cereal_tillering.jpg", units = "px", width = 900, height = 750, quality = 300)
cereal_tillering
dev.off()

jpeg("figures/cereal_rfr.jpg", units = "px", width = 900, height = 750, quality = 300)
cereal_rfr
dev.off()

jpeg("figures/cereal_height.jpg", units = "px", width = 900, height = 750, quality = 300)
cereal_height
dev.off()

jpeg("figures/cereal_top_rfr.jpg", units = "px", width = 900, height = 750, quality = 300)
cereal_top_rfr
dev.off()

jpeg("figures/cereal_coef.jpg", units = "px", width = 900, height = 750, quality = 300)
cereal_coef
dev.off()

jpeg("figures/cereal_ssr.jpg", units = "px", width = 900, height = 750, quality = 300)
cereal_ssr
dev.off()


jpeg("figures/legume_height.jpg", units = "px", width = 900, height = 750, quality = 300)
legume_height
dev.off()

jpeg("figures/legume_top_rfr.jpg", units = "px", width = 900, height = 750, quality = 300)
legume_top_rfr
dev.off()

jpeg("figures/legume_coef.jpg", units = "px", width = 900, height = 750, quality = 300)
legume_coef
dev.off()

jpeg("figures/legume_ssr.jpg", units = "px", width = 900, height = 750, quality = 300)
legume_ssr
dev.off()

jpeg("figures/LERs.jpg", units = "px", width = 900, height = 750, quality = 300)
LERs
dev.off()


jpeg("figures/repCLC.jpg", units = "px", width = 550, height = 300, quality = 300)
repCLC + theme_bw(base_size = 15)
dev.off()

jpeg("figures/repCLL.jpg", units = "px", width = 550, height = 300, quality = 300)
repCLL + theme_bw(base_size = 15)
dev.off()

jpeg("figures/repC.jpg", units = "px", width = 550, height = 300, quality = 300)
repC + theme_bw(base_size = 15)
dev.off()

jpeg("figures/repL.jpg", units = "px", width = 550, height = 300, quality = 300)
repL + theme_bw(base_size = 15)
dev.off()

jpeg("figures/repCLCHD.jpg", units = "px", width = 550, height = 300, quality = 300)
repCLCHD + theme_bw(base_size = 15)
dev.off()

jpeg("figures/repCLLHD.jpg", units = "px", width = 550, height = 300, quality = 300)
repCLLHD + theme_bw(base_size = 15)
dev.off()

jpeg("figures/repCHD.jpg", units = "px", width = 550, height = 300, quality = 300)
repCHD + theme_bw(base_size = 15)
dev.off()

jpeg("figures/repLHD.jpg", units = "px", width = 550, height = 300, quality = 300)
repLHD + theme_bw(base_size = 15)
dev.off()

## Combined figures
jpeg("figures/combined_biomass_plant.jpg", units = "px", width = 1800, height = 750, quality = 300)
cereal_biomass_plant + legume_biomass_plant + plot_annotation(tag_levels = "A")
dev.off()

jpeg("figures/combined_height.jpg", units = "px", width = 1800, height = 750, quality = 300)
cereal_height + legume_height + plot_annotation(tag_levels = "A")
dev.off()

jpeg("figures/cereal_tillering_top_rfr.jpg", units = "px", width = 1800, height = 750, quality = 300)
cereal_tillering + cereal_top_rfr + plot_annotation(tag_levels = "A")
dev.off()

jpeg("figures/combined_replication.jpg", units = "px", width = 1100, height = 1200, quality = 300)
((repCLC + theme_bw(base_size = 15)) |
  (repCLL + theme_bw(base_size = 15))) /
  ((repC + theme_bw(base_size = 15)) |
  (repL + theme_bw(base_size = 15))) /
  ((repCLCHD + theme_bw(base_size = 15)) |
  (repCLLHD + theme_bw(base_size = 15))) /
  ((repCHD + theme_bw(base_size = 15)) |
  (repLHD + theme_bw(base_size = 15))) + 
  plot_annotation(tag_levels = "A")
dev.off()

jpeg("figures/cereal_ssr_top_rfr.jpg", units = "px", width = 1800, height = 750, quality = 300)
cereal_ssr + cereal_top_rfr + plot_annotation(tag_levels = "A")
dev.off()
```
